# Benchmarking DNA Foundation Models for zero-shot variant effect prediction: the role of context, training, and architecture

**DOI:** 10.1101/2025.06.15.659748

**Authors:** Ilaria Alfisi, Francesca Ciapi, Marta Baragli, Alberto Magi

## Abstract

In this study, we systematically evaluate the performance of several DNA foundation models (NT, DNABERT, and HyenaDNA) in predicting the functional impact of genetic variants using Zero-shot scoring, a method that does not require task-specific fine-tuning. We assess the models’ sensitivity to sequence alterations introduced by Single Nucleotide Variants (SNVs), comparing their ability to capture both local and extended contextual effects. Using pathogenic, benign, and uncertain SNVs from ClinVar, we show that large multi-species NT models outperform other architectures in detecting functional consequences, not only at the mutation site but also in adjacent regions. These models exhibit superior discriminative power across variant categories, especially when aggregating Zero-shot scores over multiple surrounding tokens. Conversely, models trained solely on human sequences, such as DNABERT and HyenaDNA, show limited contextual awareness and reduced ability to differentiate variant effects. Our findings highlight the critical importance of model size, training objective, and training data diversity in shaping model performance. Furthermore, we discuss current limitations in modeling long-range dependencies in genomic sequences and suggest that innovations in transformer architectures, such as sparse attention or memory-augmented models, may provide viable paths toward scalable, genome-wide variant effect prediction.

## Introduction

The human genome harbors approximately 4 to 5 million genetic variants when compared to the reference genome [1]. These genetic variants collectively contribute to human diversity, evolution, and disease susceptibility. While the vast majority of them have little to no functional impact, a small subset can lead to significant molecular changes that alter protein function or may even result in pathogenic effects. In this context, assessing the functional impact of a variant, both within the surrounding genic region and through potential long-range regulatory effects, is crucial for distinguishing and prioritizing the few functionally relevant variants from the vast background variation, a fundamental step toward identifying disease-associated mutations.

At present, several approaches have been proposed to predict and prioritize the functional effect of genomic variants.

Some methods, like PhastCons [2], PhyloP [4], GERP [5] and SIFT [6] evaluate evolutionary conservation by measuring how conserved a genomic region remains across species, indicating its functional importance and suggesting that changes in that sequence could have deleterious effects or negatively impact the organism.

Other methods assess the impact of amino acid substitutions on protein function by combining evolutionary conservation with structural effects (e.g., PhyloP) [3] or further incorporating functional annotations (e.g., PROVEAN) [7].

In recent years, significant advances have been made in computational tools that utilize machine learning techniques such as Combined Annotation Dependent Depletion (CADD) [8], or deep neural network DeepSEA [9], that incorporate several datasets to predict the functional impact of genetic variants.

However, existing functional prediction approaches primarily focus on assessing the impact of individual variants, without considering the functional architecture of the surrounding genomic regions or evaluating their long-range effects within the broader sequence context.

The lack of methods capable of evaluating the functional effect of variants on sequence context relies on the ability of the currently available neural network framework to capture long range dependencies on sequences. Genomic sequence analysis methods predominantly relied on Convolutional Neural Networks (CNNs) and Recurrent Neural Networks (RNNs).

CNNs have been extensively utilized in genomic sequence analysis thanks to their adeptness in detecting local patterns within input sequences. Despite their prowess, CNNs faced limitations in capturing broader, long-range relationships due to inherent architectural constraints [10].

RNNs and their variants Long Short-Term Memory (LSTM), emerged as alternative tools for modelling genomic sequences, primarily due to their ability to retain temporal information. However, RNNs faced problems like vanishing and exploding gradients, which made them less effective for very long sequences [11].

The seminal paper “Attention is All You Need” by Vaswani et al. [12] marked a paradigm shift in Natural Language Processing (NLP) by introducing the transformer architecture. Unlike CNNs and RNNs, transformers leverage self-attention mechanisms to efficiently capture dependencies across an entire sequence, enabling superior performance in tasks requiring long-range contextual understanding.

In recent years, foundation models based on transformer architectures have been developed for DNA sequence analysis. The first notable transformer-based model in this domain, DNABERT [13], was adapted from the BERT architecture, originally designed for natural language processing, by tokenizing DNA sequences into k-mers, allowing it to capture contextual relationships within genomic data.

More recently, the Nucleotide Transformer (NT) [14] has emerged as a more advanced foundation model, leveraging the transformer architecture for comprehensive DNA sequence analysis. NT builds on DNABERT’s foundational concepts, incorporating a larger training dataset and more sophisticated training techniques to enhance performance.

Alongside transformer-based models, alternative foundation models have also been introduced, such as HyenaDNA [15], which takes a fundamentally different approach to long-range genomic sequence modeling. Unlike transformers, which rely on self-attention, HyenaDNA uses a Structured State-Space Model (SSM) to efficiently capture long-range dependencies at single-nucleotide resolution significantly reducing memory and computational costs.

These models have been applied to promoter prediction, transcription factor binding site identification, splice site detection, enhancer sequence prediction, and prioritization of functional genetic variants.

In this study, we investigated the potential of three DNA foundation models (NT, DNABERT, and HYENA) without any task-specific fine-tuning to evaluate the functional impact of SNVs and assess how these variants influence the surrounding sequence context.

To this end, we analysed sequences containing SNVs from various functional classes, annotated in the ClinVar database as pathogenic, uncertain, or benign. We show that DNA foundation models differ significantly in their ability to predict Single Nucleotide Variant (SNV) impact and capture the broader sequence context, depending on factors such as model architecture, training objective, parameter count, and the diversity of the training dataset. The findings of this study offer valuable guidance for the development and selection of future DNA foundation models in the context of functional variant interpretation.

## Results

### Zero-shot scores of large DNA model capture both the local variant effect and the impact on surrounding context

DNABERT, NT and HyenaDNA employ different training strategies to capture meaningful sequence representations. DNABERT and NT follow the Masked Language Model (MLM) approach, where random k-mers in a DNA sequence are masked, and the model learns to predict them based on surrounding context (see Table 1).

**Table 1:** Architectural and training details of the DNA language models benchmarked in this study. Models vary in total number of parameters, number of layers, embedding dimensions, input token lengths, objectives function (MLM = Masked Language Modeling; NTP = Next Token Prediction), and training datasets. “NT” models are part of the Nucleotide Transformer family and differ by number of parameters and training dataset (multispecies, 1000 Genomes, or human reference, see *Methods* for more details). All models use k-mers as input tokens, with k ranging from 1 to 6 depending on architecture.

| Model | Parameters | Layers | Embedding | Token | Objective Function | Dataset | K-mers |
| --- | --- | --- | --- | --- | --- | --- | --- |
| NT_2.5B | 2.5B | 32 | 2560 | 1000 | MLM | Multispecies | 6 |
| NT_1G_2.5B | 2.5B | 32 | 2560 | 1000 | MLM | 1000G | 6 |
| NT_1G_500M | 500M | 24 | 1536 | 1000 | MLM | 1000G | 6 |
| NT_HR_500M | 500M | 24 | 1536 | 1000 | MLM | Human Reference | 6 |
| NT_500M | 500M | 29 | 1536 | 1000 | MLM | Multispecies | 6 |
| NT_500M | 500M | 24 | 1024 | 1000 | MLM | Multispecies | 6 |
| NT_100M | 100M | 22 | 768 | 1000 | MLM | Multispecies | 6 |
| DNABERT | 110M | 12 | 768 | 512 | MLM | Human Reference | 6 (overlapping) |
| Hyena_1 | 436k | 2 | 128 | 1025 | NTP | Human Reference | 1 |
| Hyena_32 | 1.6M | 4 | 128 | 6000 | NTP | Human Reference | 1 |

On the other hand, HyenaDNA, utilizes Next Token Prediction (NTP), training the model to predict the next nucleotide given the preceding sequence (*see Methods*).

Moreover, while DNABERT and NT rely on k-mers of 6 base pairs (bp) for tokenization, HyenaDNA employs a single-base tokenizer, maintaining nucleotide-level specificity (see Table 1 and *Methods* for more details). These distinct strategies reflect different trade-offs: MLM-based models excel in capturing bidirectional context and genomic motifs, whereas HyenaDNA is optimized for processing extremely long sequences, leveraging efficient convolutions instead of attention mechanisms.

For all the three models, the learned representations are encoded by the embeddings from the last layers of the model that capture the model’s understanding of the entire input sequence, enabling the extraction of recurring patterns and relationships between nucleotides or functional regions.

Consequently, embeddings can be used to measure the similarity between DNA sequences, helping to determine how similar two sequences are in terms of meaning.

The embeddings are then passed through a fully connected layer (or “linear layer”), which is a weight matrix learned during the model’s training. The output of this operation is a vector of logits, representing unnormalized scores for each class. These logits are often then passed through a softmax function to obtain the probability that a nucleic acid k-mer occurs at a position in a DNA sequence given the surrounding context.

This intrinsic property of foundation DNA models can be used to score sequence variations by comparing the embeddings (or prob-logit distributions) assigned to the mutated k-mer with those assigned to the reference k-mer.

Evaluating the differences between the embedding layers or the probability distributions of two sequences, one carrying the mutated allele and a second carrying the reference allele, can be thus used to evaluate the functional impact of a variant.

Moreover, since language models can capture long-range interactions within a sequence, differences between embeddings or probabilities can also measure the effect of a variant on all other tokens in the context.

The differences between embeddings or probabilities of two sequences, can be evaluated by using various distance or similarity, and these measures are defined as Zero-shot scores since they are used to predict classes that were not seen by the model during training.

In this work we used, as Zero-shot-based scores, the Cosine Similarity, the Manhattan and Euclidean distances of the embeddings and the Hellinger distance, the Jensen-Shannon divergence and the Cross-Entropy for the probability distributions (*see Methods* and Figure 1).

**Figure 1:**
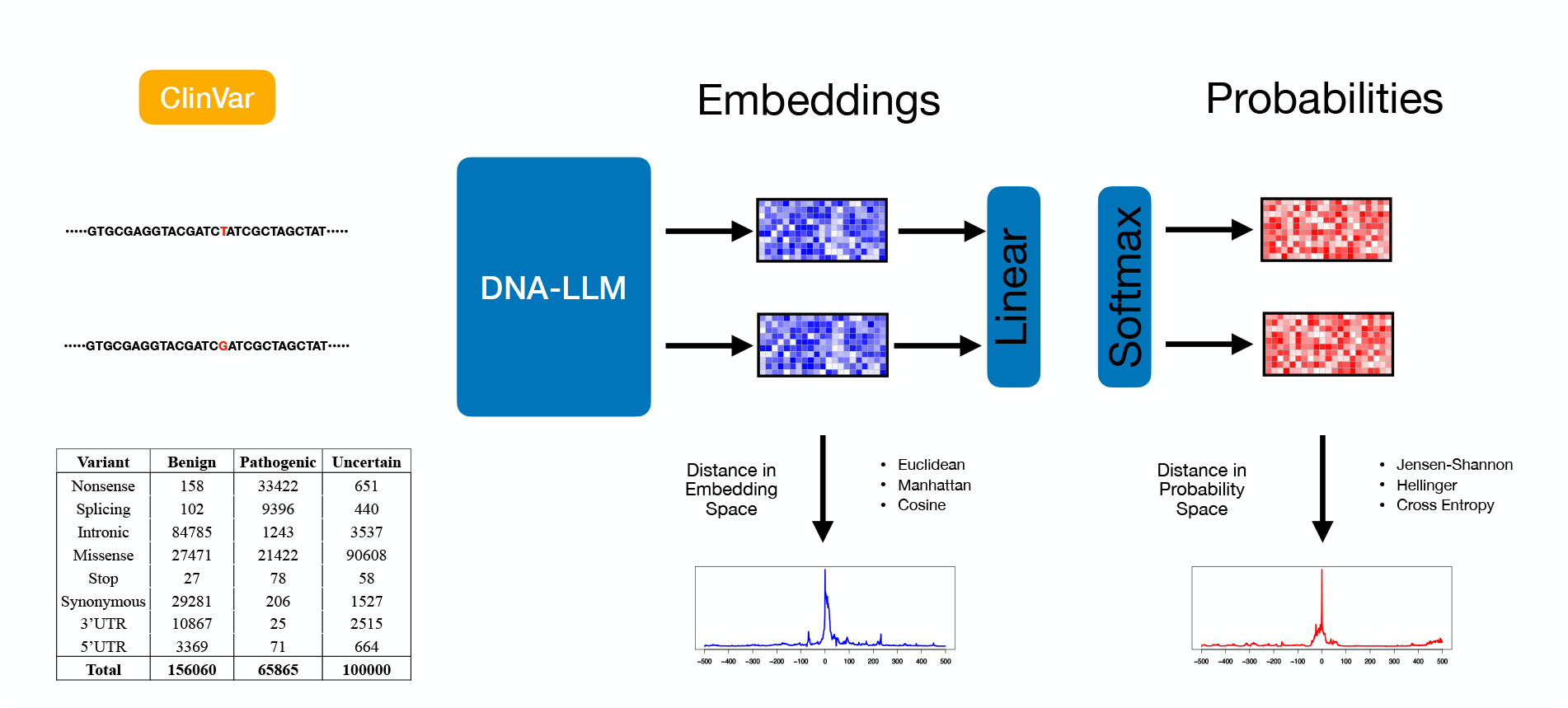
Schematic overview of the Zero-shot scoring procedure used to compare DNA foundation models on ClinVar SNVs. For each SNV, a DNA sequence centered on the variant (6 Kb for NT models, 512 bp for DNABERT, 1000 bp for HYENA 1) was extracted from the human reference genome (GRCh38/hg38), and two sequences were created: one carrying the reference allele and one the ClinVar allele. Each sequence pairs were processed through all ten foundation models to obtain both the final-layer embeddings and the output probabilities (via a linear layer followed by softmax). Embedding vectors were used to compute Cosine Similarity, Manhattan, and Euclidean distances, while probability distributions were compared using Hellinger distance, Jensen-Shannon divergence, and Cross-Entropy.

To compare the performance of the ten transformer models (seven NT, DNABERT and two HYENA) we used 320,000 SNVs selected among benign, pathogenic and uncertain significance classification schemes of the ClinVar database (*see Methods*).

In particular, we used 156060 Bening SNVs, 65865 Pathogenic and 108386 of uncertain significance that belong to eight distinct functional classes: nonsense, splicing, stop, missense, synonymous, intronic, 3’UTR and 5’UTR (see table Figure 1).

The great majority of Benign SNVs were intronic or synonymous (73%), pathogenic were mainly nonsense and missense (85%), while uncertain were almost exclusively missense (90%). For each variant of our datasets, from the human reference genome (GRCh38/hg38) we extracted a 6 Kb DNA segment centered at the SNV position and we then created two sequences, one carrying the reference allele and a second carrying the ClinVar allele at the central position (for DNABERT we selected 512 bp sequence, while for HYENA 1 1025 bp sequence, see Table 1).

All sequences were then subjected to inference for each of the ten models (Figure 1), and for each pair of sequences (reference vs Clinvar mutation), we calculated the three Zero-shot scores in the embedding space (Cosine Similarity, Manhattan and Euclidean distances) and the three in the probability space (the Hellinger distance, the Jensen-Shannon divergence and the Cross-Entropy).

With this procedure we are able to obtain a similarity/distance measure for each token pair between the reference sequence and the sequence with the ClinVar mutation, allowing us to assess not only the local impact of the mutation but also its effect across the entire sequence context.

Figure 2 and supplemental Figures S1–S14 report the six similarity/distance profiles (Cosine Similarity, Manhattan, Euclidean, Jensen-Shannon and Cross-Entropy.) averaged across the eight distinct functional classes for SNVs belonging to Benign, Pathogenic and Uncertain categories.

**Figure 2:**
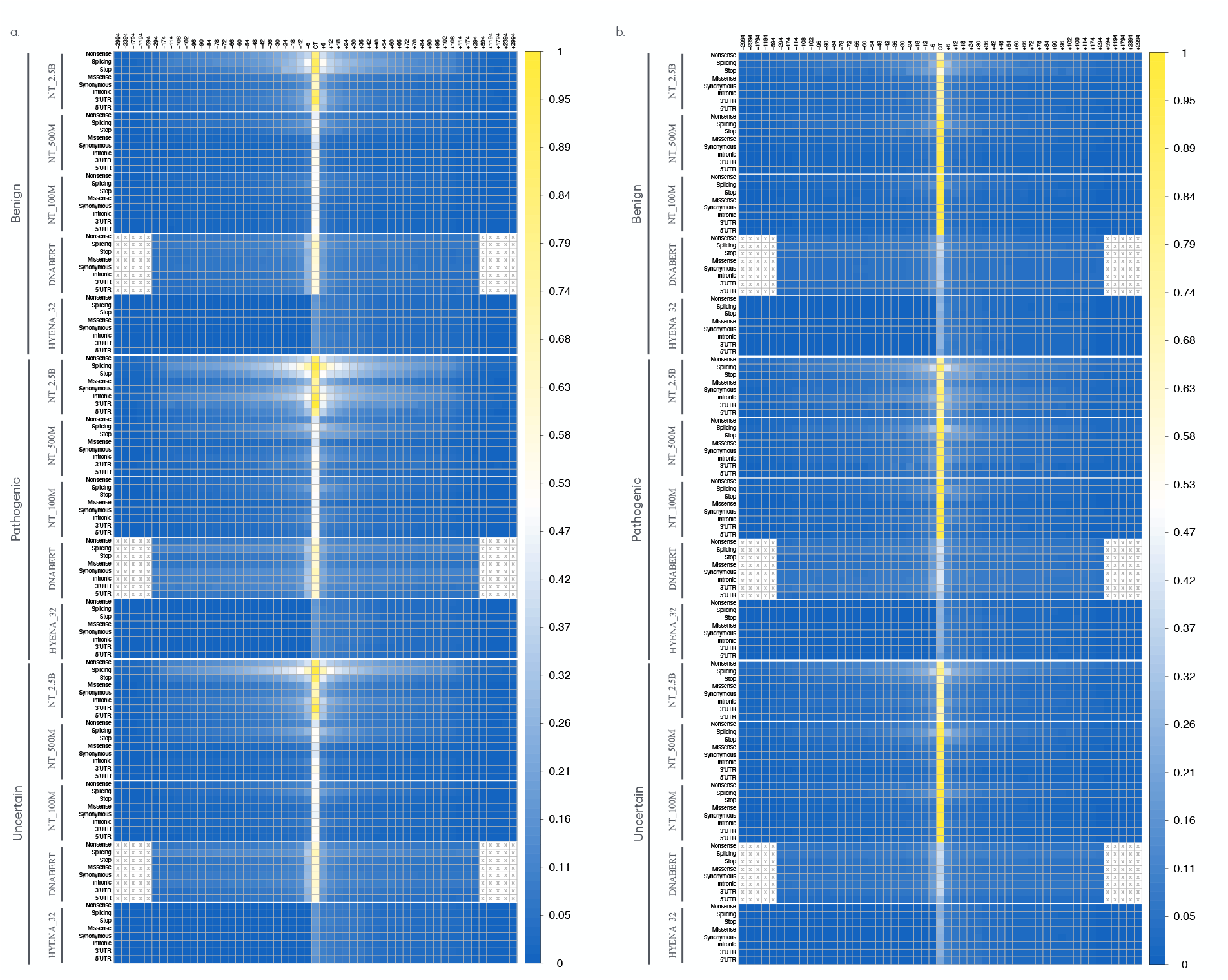
Figure displays the Zero-shot score profiles across all tokens of the sequences analyzed using the DNA foundation models. Each row in the heatmap shows the trend of Zero-shot score profiles calculated for sequence pairs (with ClinVar and reference bases) by averaging, for each token in the sequence, the Zero-shot scores obtained by different models for each genetic consequence category (nonsense, stop, splicing, missense, synonymous, intronic, 5’UTR, and 3’UTR). The results are presented by distinguishing the three clinical impact categories (Benign, Pathogenic, and Uncertain) for the five main foundation models used (NT_2.5, NT_500, NT_100, DNABERT, and HYENA_32). The intensity of each cell reflects the value of the Zero-shot score according to the color legend on the right. Values are normalized by dividing each score by its maximum value. Panel (a) shows the Zero-shot score profiles based on Euclidean distance in the embedding space, while panel (b) shows those based on Jensen-Shannon divergence in the probability space.

For all the models, the maximum distance (i.e., minimum similarity) occurs at the central token of the sequence (the token carrying the ClinVar mutation), both in the embedding and probability spaces. The highest distances are observed in the multi-species NT models, followed by the NT models trained exclusively on human sequences, and then by the DNABERT and HyenaDNA models.

Remarkably, in the embedding space of multi-species NT models, the lowest distances are observed for missense and synonymous variants.

A more in-depth analysis of these images also reveals that the similarity/distance profiles of NT multi-species models with a high number of parameters (mainly NT_2.5B, NT_500M, NT_250M) not only show a peak at the central token (the one with the ClinVar mutation), but also exhibit deviations in the adjacent tokens. The extent of these deviations (i.e., the number of neighboring tokens showing noticeable shifts in similarity or distance values) correlates with functional impact (in stop, splice, and nonsense variants, the effect propagates across more tokens than in synonymous, missense, intronic, and UTR variants) and the ClinVar category of the mutations (pathogenic variants show more deviations than Uncertain, which in turn show more than Benign).

Surprisingly, while benign variants with low functional impact (synonymous, missense, intronic, UTR) show deviations localized to just the central token (or a few adjacent tokens), pathogenic variants with the same functional impact exhibit similarity/distance profiles with deviations extending across a larger number of tokens.

Conversely, NT models with a low number of parameters (NT_100M) and those trained only on human sequences (NT_1G and NT_HR) fail to capture the effect of the variants on the adjacent tokens as described above.

Regarding the other models, while DNABERT cannot assess the impact of a variant on adjacent tokens, both HyenaDNA (see Figure S13) models are capable of capturing long-range functional effects, although no significant differences are observed between the different types of variants.

Additionally, while in the NT and DNABERT models the effect of the variant is observed in the surrounding tokens, both in the preceding direction (to the left) and the succeeding direction (to the right), in the HyenaDNA models the effect is detected only in the succeeding tokens. This discrepancy can be ascribed to the different training methods used: MLM for DNABERT and NT, and NTP for HYENA. In MLM-based models, the model learns to predict masked tokens from both directions, allowing bidirectional context capture. In contrast, the NTP approach, used in HYENA, predicts only the next token in a sequence, which restricts the variant’s effect to the succeeding tokens.

Finally, we observed notable distinctions between probability and embedding spaces. The embedding representations, across all three similarity measures employed, demonstrate a superior ability to capture long-range functional effects on adjacent tokens. In contrast, the probability space is considerably less effective, particularly when using Cross-Entropy and Hellinger distance, which fails to capture the effect of the ClinVar token on adjacent tokens (Supplemental Figure S1–S14).

These findings demonstrate that very large NT models, especially in embedding space, can evaluate not only the local effect of a genomic variant but also its impact within the surrounding sequence context.

### Zero-shot scores of the mutated token distinguish high- vs low-impact variants, but not pathogenic from benign ones

In order to quantify the capability of the ten DNA-models to discriminate variants with different genetic consequences, we compared the distribution of the Zero-shot score between each pair of the eight distinct genetic variants classes.

To this end, we first used the similarity/distance measure of the token carrying the ClinVar variant and for each pair of genetic variants classes we calculated the Wilcoxon U-statistic.

The Wilcoxon U-statistic (*see Methods*) compares the ranks of data points from two independent groups and tells us how many times a data point from one group precedes a data point from the other group. A smaller U statistic suggests that values in one group are generally lower than the other.

Figure 3 and supplemental Figure S15 clearly show that the ten DNA-models have very different discriminative abilities, which are closely dependent on the number of parameters and on the space, either embedding or probability, where the Zero-shot scores are computed.

**Figure 3:**
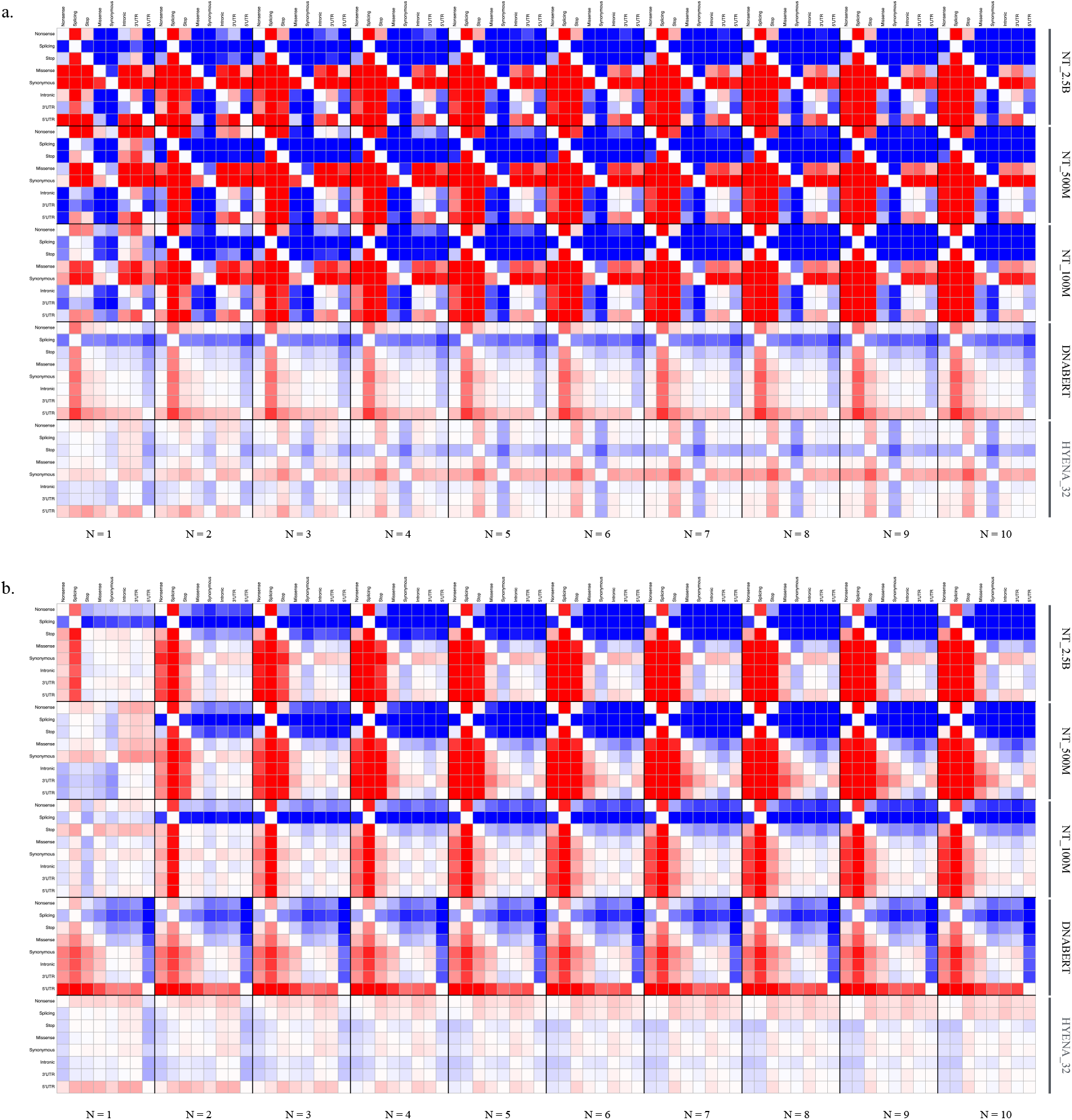
The figure illustrates the ability of the five main foundation models used (NT_2.5B, NT_500M, NT_100M, DNABERT, and HYENA_32) to discriminate variants with different genetic consequences. Each cell in the color plot represents the normalized Wilcoxon U-statistic, comparing the distribution of the CCS-N, obtained by summing the Zero-shot scores of the N (N=1, only the central token) tokens closest to the central token carrying the ClinVar variants, between two classes of genetic variants. For each pair of variant types, the scores were compared across all three clinical impact categories (Benign, Pathogenic, and Uncertain). The intensity of each cell reflects the normalized Wilcoxon U-statistic value, as indicated by the color legend on the right. Panel (a) shows the results for Euclidean distance calculated in the embedding space, while panel (b) for Jensen-Shannon divergence in the probability space.

In general, the best performance in discriminating between genetic variant classes (high U-statistic values) is achieved by Zero-shot scores calculated in the embedding space of all models, especially for multi-species NT models with a large number of parameters (NT_2.5B, NT_500M, Figure 3.a and Supplemental Figure S15.a-c), and this performance decreases as the number of model parameters decreases.

Specifically, multi-species NT_2.5B embeddings demonstrate strong discriminative power (high U-statistic values), effectively distinguishing missense, synonymous, and 5’UTR variants from all other genetic categories. Conversely, NT_500M, NT_250M, and NT_100M models primarily differentiate missense, synonymous, and nonsense variants from the rest.

Interestingly, across all multi-species NT models, intronic, 3’UTR, and 5’UTR variants exhibit higher Zero-shot scores than missense and synonymous variants, consistent with the patterns observed in the profile plots (see Figure 3).

By contrast, all NT models trained exclusively on human sequences, as well as DNABERT and HyenaDNA models, exhibit significantly lower discriminative power (see Figure 3.a and Supplemental Figure S15.a-c).

Notably, no substantial differences were observed among the three Zero-shot scores computed in the embedding space. On the other hand, Zero-shot scores computed in the probability space vary significantly in their discriminative power across different models (see Figure 3.b and Supplemental Figure S15.d-f). For both Jensen and Cross-Entropy scores, DNABERT performs remarkably well, achieving high U-statistic values in distinguishing variants with strong functional impact (stop, splicing, nonsense, and missense) from those with lower impact (synonymous, intronic, 5’UTR, and 3’UTR). Similarly, with these two metrics, the NT_2.5B model effectively separates nonsense and splicing variants from all others, whereas all other NT models and HyenaDNA models yield poor results (Figure 3.b and Supplemental Figure S15.d-f).

Regarding the Hellinger metric, all multi-species NT models outperform the other two probability-based scores, with NT_2.5B standing out as the most effective in differentiating high-impact from low-impact variants. In contrast, all other models exhibit poor performance (see Supplemental Figure S15.d-f).

As a further step, we decided to evaluate the capability of six Zero-shot scoring methods to distinguish between pathogenic and benign variants within the same functional category.

To this end, we computed the Wilcoxon U-statistic to compare the score distributions of pathogenic and benign variants across each functional category.

The results presented in Figure 4 and Supplemental Figure S16 indicate that none of the DNA-based models, whether evaluated using distances in embedding space or probability space, achieve a Wilcoxon U-statistic above 0.6. The only exceptions are the multi-species NT models (NT_2.5B, NT_500M, and NT_250M), which achieve U-statistics greater than 0.7 for synonymous variants in the embedding space (Figure 4.a and supplemental Figure S16.a-c).

**Figure 4:**
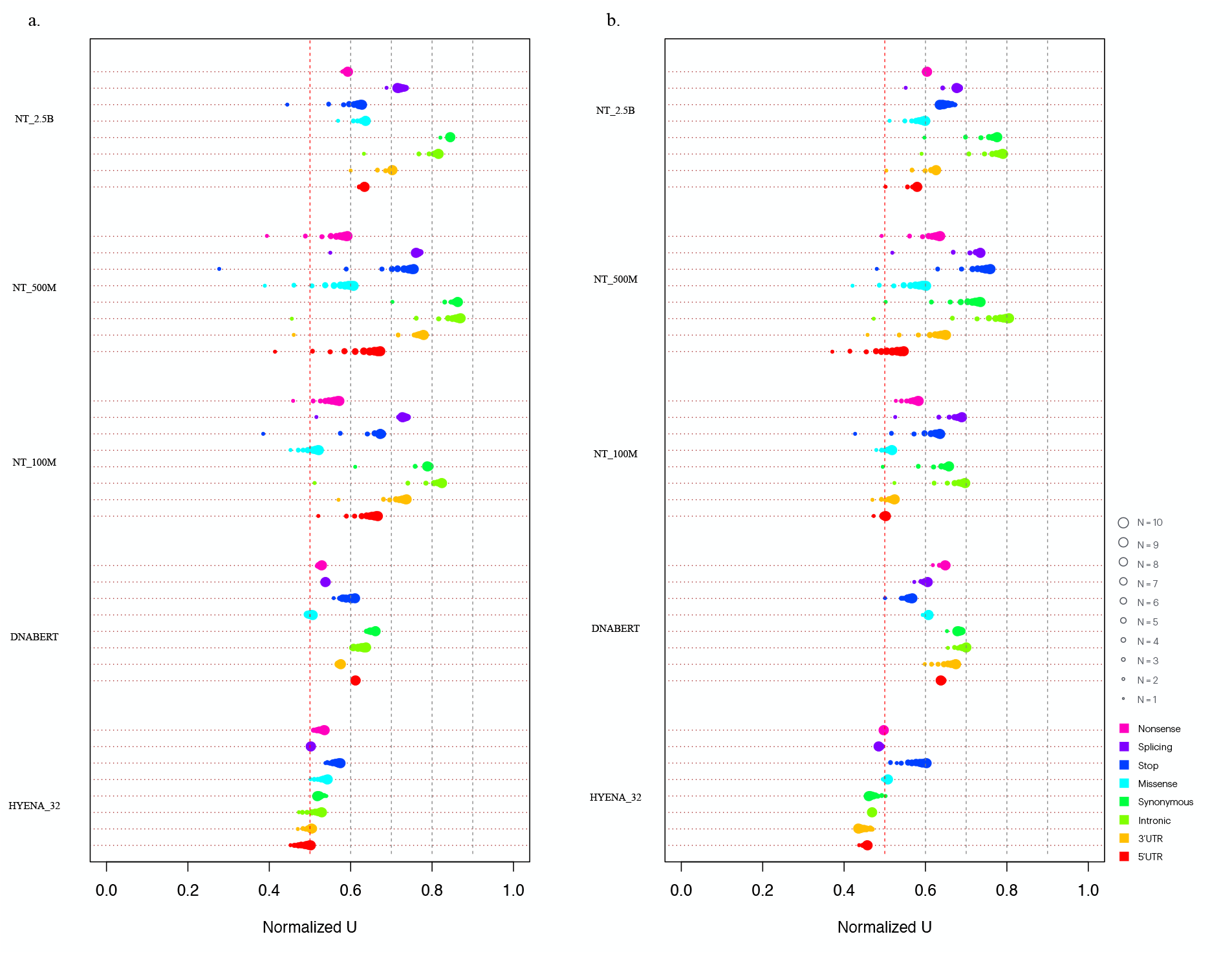
The figure illustrates the ability of the five DNA foundation models to distinguish benign from pathogenic variants. Each point in the dot chart represents the normalized Wilcoxon U-statistic, comparing the distribution of CCS-N, obtained by summing the Zero-shot scores of the N tokens closest to the central token carrying the ClinVar variants, between benign and pathogenic variants within the same genetic consequence class. Panel (a) shows the results for Euclidean distance calculated in the embedding space, while panel (b) for Jensen-Shannon divergence in the probability space.

Unexpectedly, the NT_500M, NT_250M, and NT_100M models yield U-statistic values below 0.5 for nearly all variant types across most Zero-shot scoring methods (except for Hellinger, Supplemental Figure S16). This suggests that these models assign higher scores to benign variants than to pathogenic ones, failing to capture the functional context in which the variants occur.

Conversely, while DNABERT does not achieve good results, it maintains U-statistic values above 0.5 for almost all variant classes. This indicates that both embeddings and probability-based scores allow, to some extent, the distinction between pathogenic and benign sequences.

Consistent with previous findings on functional category discrimination, the HyenaDNA and NT models trained exclusively on human sequences perform the worst. DNABERT, on the other hand, continues to achieve better results with Zero-shot scores in probability space than in embedding space, although it never exceeds *U >* 0.6 in any analysis. Still, DNABERT consistently maintains U-statistic values above 0.5 across nearly all variant classes, reinforcing the notion that its embeddings and probability-based scores provide some capacity to differentiate pathogenic from benign sequences. As observed in earlier analyses, while Zero-shot scores computed in probability space exhibit considerable variation in predictive performance, the three embedding-based scores show no substantial differences.

### Combining Zero-shot scores across multiple tokens improves DNA models’ ability to distinguish high- vs low-impact and pathogenic vs benign variants

The analysis of the Zero-shot-score profiles (Figure 2 and supplemental Figures S1–S14) shows that changing the central token not only affects the score of that token but also impacts the scores of the neighboring tokens, especially for variants with higher genetic consequences, suggesting that including more tokens in predicting the functional impact of a variant could increase its predictive value.

For this reason, we decided to evaluate the discriminative power of the Cumulative Context Score (CCS-N) as the sum of the Zero-shot score at the central variant-associated token and the Zero-shot scores of the N closest surrounding tokens, added incrementally in order of proximity (with N that ranges between 1 and 10, *see Methods*). As in the previous section, we first evaluated the capability of the six Zero-shot scores to discriminate variants with different genetic consequences, by comparing the distribution of the CCS-N between each pair of the eight distinct genetic variants classes by using the Wilcoxon U-statistic. The results reported in Figure 3 and supplemental Figure S15 show that CCS-N increase the discriminative ability of all models, particularly multi-species NT models with a high number of parameters, both in the embedding space and in the probability space. Similarly to what was observed in the central token analysis, CCS-N in the embedding space show no significant differences among the three Zero-shot scoring methods. Summing scores across multiple tokens progressively enhances the ability to distinguish high-impact variants (nonsense, splicing, and stop) from low-impact ones (missense, synonymous, intronic, 3’UTR and 5’UTR, Figure 3 and supplemental Figure S15). Discriminative performance stabilizes with 3-4 tokens. As expected, NT models trained on human sequences, DNABERT, and HyenaDNA models exhibit only marginal improvements when using CCS-N in the embedding space (Figure 3.a and supplemental Figure S15.a-c). In probability space, the models that benefit the most from CCS-N are the multi-species NT_500M, NT_250M, and NT_100M models, whereas all other models show only minor improvements (Figure 3.b and supplemental Figure S15.d-f).

A particularly important observation is that, across all multi-species NT models, CCS-N in probability space effectively discriminate nonsense, splicing, stop, missense, and synonymous variants from all others, especially with Jensen and Hellinger measures (Supplemental Figure S15 d-f). This suggests that, in probability space, missense and synonymous variants receive higher scores compared to embedding space. Similarly to the embedding space, the probability space shows only marginal improvements for NT models trained on human sequences, DNABERT, and HyenaDNA when using CCS-N. Overall, in probability space, the best discriminative performance is achieved by the multi-species NT_500M model using the Hellinger distance, followed by the other multi-species models (NT_2.5B, NT_250M), which also perform well in embedding space. As a next step, we assessed the ability of the CCS-N to distinguish between benign and pathogenic variants with the same genetic consequence.

Consistent with the previous analysis, using the CCS-N enhances the ability of all models to discriminate pathogenic from benign variants, particularly high-parameter multi-species NT models in embedding space (Figure 4 and supplemental Figure S16). In this setting (Figure 4.a and supplemental Figure S16.a-c), the NT_500M model effectively differentiates pathogenic from benign variants, achieving U-statistics between 0.7 and 0.8 for splicing, stop, and 3’UTR variants, and between 0.8 and 0.9 for synonymous and intronic variants. The NT_2.5B model yields similar results, except for stop variants.

In contrast, CCS-N computed in probability space exhibit weaker performance, with U-statistics exceeding 0.7 only for intronic and synonymous variants in the NT_2.5B model (using Jensen-Shannon, Figure 4.b and supplemental Figures S16.d-f) and for intronic, synonymous, stop, and splicing variants in NT_500M (using Hellinger and Jensen-Shannon distances). All other models, including NT models trained on human sequences, DNABERT, and HYENA, show only marginal improvements in discriminative ability.

### DNA foundation models have learned the structural patterns of both coding and non-coding regions of the genome

Language Model (LM) based on Transformers estimate the probability of a token given its context, and for this reason, the Zero-shot score profiles should measure the impact of a variant not only locally but also within the broader context, distinguishing between functionally relevant genomic segments and those with a lower functional impact. In this scenario, variants with a more severe genetic consequence (splicing, nonsense, and stop) should more significantly alter the Zero-shot scores across the entire context compared to variants with a less severe genetic consequence (synonymous, missense, and intronic). Additionally, if the models have learned the functional structure of the genome, this impact should be greater in coding elements (exons) compared to non-coding elements (introns and intergenic regions).

To test this hypothesis for each variant we calculated the normalized Mann-Whitney U statistic between the distribution of Zero-shot-scores in coding vs non-coding tokens (*see Methods*). If the normalized Mann-Whitney U statistic is greater than 0.5, it indicates that the Zero-shot scores in coding regions are higher than in non-coding regions. This suggests that the foundation models have successfully learned the functional structure of the genome.

As in all other analyses conducted in this study, the results obtained with the Zero-shot scores calculated in the embedding space are superior to those in the probability space, achieving higher normalized U-statistic values across all models (Figure 5 and Supplemental Figure S17).

**Figure 5:**
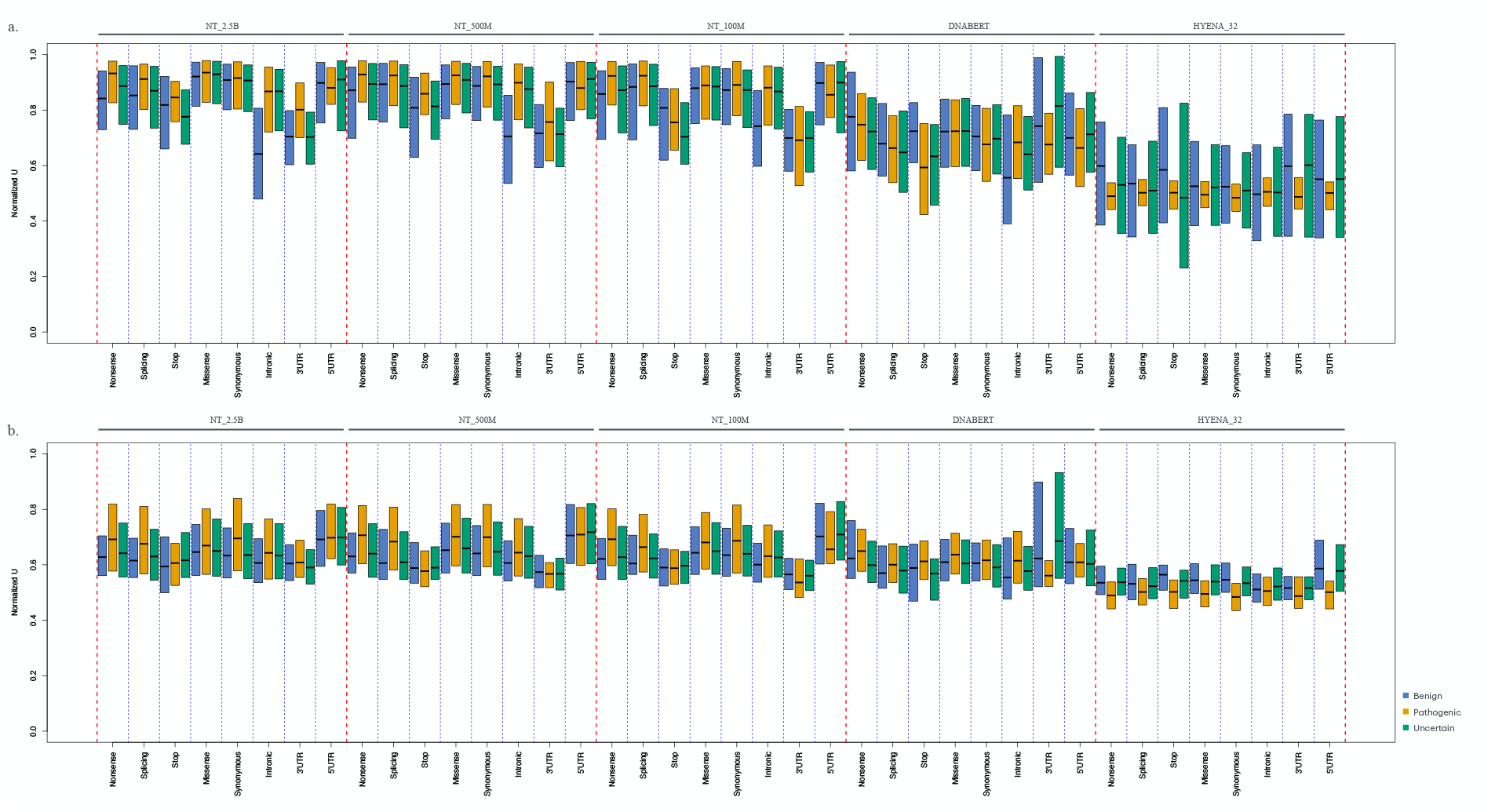
The figure illustrates the ability of the five foundation models to assess the impact of ClinVar variants on the functional structure of the genome. The box plots show, for each genetic consequence class, the distribution of normalized U-statistics comparing Zero-shot scores between coding elements (exons) and non-coding elements (introns). Panel (a) shows the results for Euclidean distance calculated in the embedding space, while panel (b) for Jensen-Shannon divergence in the probability space.

In accordance with the results found in previous analyses Figure 5 and supplemental Figure S17 show that multi-species NT models (from NT_2.5B to NT_100M) obtain the highest U statistic for Zero-shot-scores in both embedding and probability spaces (with the exception of NT_2.5B with Hellinger distance), suggesting that these models are the ones that have best learned the functional structure of the genome. However, although embedding similarity, on average, achieves a higher U-statistic value and thus a greater ability to discriminate the impact of a variant on elements with higher genomic functionality, the Jensen and Cross-Entropy scores in the probability space are more correlated with the functional impact of the variant. They achieve higher U-statistic values for high-impact variants (nonsense and splice) and lower values for low-impact variants (synonymous, 5’UTR, and introns).

As expected, HYENA, DNABERT, and human-based NT models achieve the worst results also in these analyses, suggesting that these are the models that have learned much less about the functional structure of the genome.

## Discussion and Conclusion

In this study, we evaluated the performance of various DNA foundation models (NT, DNABERT, and HYENA) in assessing the functional impact of genetic variants using Zero-shot scoring methods, an approach that computes scores directly from the pre-trained model without any task-specific fine-tuning. Until now, Zero-shot scoring methods have mainly been used to predict the functional impact of variants at a global scale, without evaluating how DNA foundation models capture the influence of a variant within its broader sequence context. In this study, we compare Zero-shot scores from three classes of DNA foundation models, assessing not only their ability to prioritize variants by functional impact, but also their capacity to model the sequence context in which these variants have the strongest effects.

Zero-shot scores are derived by comparing the differences in embeddings or probability distributions between the tokens of the reference and mutated sequences, thereby reflecting the model’s inherent ability to capture the functional consequences of genetic alterations.

To this end we used variants with different functional impact selected among benign, pathogenic and uncertain SNVs from the ClinaVar database.

Across all models, the central token (the one directly affected by a SNV) exhibited the largest deviation (i.e., lowest similarity or highest distance) between the mutated and reference sequences, confirming that these models are sensitive to local changes. Notably, multi-species NT models (especially those with large numbers of parameters e.g., NT_2.5B and NT_500M) generated the highest distances in both embedding and probability spaces. These models not only detected the local effect of the variant but also exhibited alterations in adjacent tokens. For variants with higher predicted functional impacts (e.g., stop, splice, and nonsense), the perturbation propagated over several neighboring tokens, while for variants with lower functional impact (such as synonymous or certain missense variants) we observed a much more confined effect.

In contrast, DNABERT and NT models trained exclusively on human sequences (NT_1G, NT_HR) demonstrated an impaired ability to capture these adjacent token effects, while HyenaDNA models, owing to their NTP training, captured alterations only in the succeeding tokens, thus missing bidirectional contextual cues. These observations suggest that the richness of the training data (multi-species versus human-only) and model dimension and architecture play a critical role in capturing not just the local variant effect but also its influence on the broader sequence context.

Similar results were observed when we evaluated the ability of the DNA foundation models to discriminate between genetic variants with different functional consequences using Zero-shot scores computed for the central token. As expected, the models’ performance is strongly influenced by both the number of parameters and the feature space in which the scores are computed. The best discrimination between variant classes was achieved by the embeddings of large multi-species NT models (e.g., NT_2.5B and NT_500M) that demonstrated robust capacity to differentiate variants with high functional impacts (such as non-sense, splicing, and stop variants) from those with lower impact (including missense, synonymous, and UTR variants).

In contrast, NT models trained solely on human sequences, along with DNABERT and HYENA, exhibited markedly lower discriminative power in the embedding space.

Interestingly, when scores were computed in the probability space, the discriminative abilities varied considerably among models. Both Jensen-Shannon and Cross-Entropy metrics allowed DNABERT to achieve relatively good discriminative performance separating variants with strong functional impact from those with weaker effects.

None of the models’ central token Zero-shot scores were able to discriminate between benign and pathogenic variants within the same functional category, irrespective of whether the scores were derived from the embedding or probability space.

Surprisingly, multi-species NT models (specifically NT_500M, NT_250M, and NT_100M) yielded U-statistic values below 0.5 across nearly all variant types, implying that these models sometimes assigned higher scores to benign variants than to pathogenic ones.

Recognizing that the effect of a variant might extend beyond the central token, we extended our analysis by summing the Zero-shot scores across the N closest tokens (the “N-sum” approach). This multi-token analysis significantly enhanced the discriminative power of the models. For multi-species NT models with high parameter counts, the CCS-N (with optimal performance reached using 3-4 tokens) allowed for a more refined separation between high-impact variants (e.g., nonsense, splicing, stop) and lower-impact ones. In embedding space, the improvement was consistent across the three Zero-shot scoring methods, indicating that aggregating scores over multiple tokens better captures the extended context of variant effects. In probability space, a similar trend was observed, particularly for NT_500M, NT_250M, and NT_100M models, although the improvement was more modest for models trained solely on human sequences, as well as for DNABERT and HYENA.

In addition to differentiating functional classes, we further assessed the ability of the CCS-N to discriminate between benign and pathogenic variants within the same functional category. Aggregating scores over adjacent tokens markedly enhanced this differentiation, with the strongest performance again observed in high-parameter, multi-species NT models in the embedding space. For example, the NT_500M model achieved U-statistics between 0.7 and 0.8 for splicing, stop, and 3’UTR variants, and between 0.8 and 0.9 for synonymous and intronic variants, while NT_2.5B yielded similar results except for stop variants. In contrast, models computed in the probability space, along with NT models trained exclusively on human sequences, DNABERT, and HYENA, showed only modest improvements.

Overall, the multi-token approach reinforced the notion that a variant’s impact is not isolated to its immediate location, and that leveraging adjacent contextual information can substantially increase predictive accuracy.

As a final step, to assess whether DNA foundation models have captured the functional structure of the genome, we compared Zero-shot scores between coding (exonic) and non-coding (intronic/intergenic) regions across all variant classes. Once again, the multi-species NT models with a high number of parameters performed best, suggesting they have more effectively learned the genome’s functional architecture, producing higher scores in coding elements and lower scores in non-coding regions. In contrast, DNABERT, HYENA, and NT models trained only on the human genome consistently showed inferior performance.

DNABERT is a Transformer-based model with an architecture closely resembling that of NT models and, in principle, should have learned the functional structure of the genome. However, our analyses reveal substantial limitations: DNABERT performs poorly in distinguishing variants with different functional impacts and in evaluating their long-range effects on the surrounding sequence context. These short-comings are likely due to its relatively small model size (12 layers, 768 embedding dimensions, and 12 attention heads per layer), the restricted input sequence length (up to 512 base pairs), and the fact that it was trained solely on human genome sequences.

Recently, the authors of DNABERT introduced and published an updated foundation model, DNABERT2, in which they replaced k-mer tokenization with Byte Pair Encoding (BPE), a statistics-based data compression algorithm that constructs tokens by iteratively merging the most frequent co-occurring genomic segments in the corpus.

Unfortunately, DNABERT2 cannot be included in the type of analysis we performed, due to the inability to compute Zero-shot scores between the tokens of the reference and mutated sequences. This is because a single base change in the mutated sequence may alter both the content and number of resulting tokens, making direct comparison of the two sequences infeasible.

Regarding HyenaDNA models, these consistently showed the weakest performance, both in distinguishing variants with different functional impacts and in assessing their long-range influence across the full sequence context.

As also noted by the original authors [15], HyenaDNA is significantly smaller in size (7 million parameters) compared to other DNA foundation models, and it was trained exclusively on the human genome. This limited training diversity likely reduces the model’s ability to capture conserved regulatory signals across species, signals that are often essential for accurately predicting the effects of SNVs at a distance.

While expanding the training dataset to include DNA sequences from other species and increasing model size may improve long-range modeling capabilities, we believe the primary limitation of HyenaDNA lies in its training objective. Despite being designed for efficient processing of long contexts, HyenaDNA was trained using a next-token prediction strategy. This approach emphasizes linear sequence modeling and local dependencies, but it is inherently less effective at learning non-local interactions. In contrast, MLM requires the model to integrate global sequence context in order to reconstruct masked elements, encouraging a richer understanding of the overall structure.

As a result, HyenaDNA can process long sequences, but may fail to represent them with sufficient granularity and discrimination. Relevant signals may be compressed or diluted over extended contexts, reducing the model’s ability to capture the localized effect of a single variant.

Our comprehensive analysis shows that NT models possess unique strengths in predicting the functional impact of SNVs. While all models are sensitive to local sequence alterations, as reflected by Zero-shot scores at the central token, only the embeddings from large, multi-species NT models consistently capture broader contextual effects. By aggregating scores across multiple tokens (the CCS-N approach), we further improve variant discrimination, particularly for high-impact mutations that affect sequence context beyond the immediate site. These results suggest that such models have more effectively learned the functional architecture of the human genome and are better equipped to predict the effects of genomic variants. Their superior performance likely stems not only from architectural choices (such as rotary position embedding, gated linear units, and a higher parameter count) but also from the inclusion of sequences from multiple species in the training dataset.

Taken together, the results and analyses presented in this study highlight the key factors necessary for developing DNA foundation models capable of capturing the complex functional architecture of genomes. These include: the training objective, where MLM proves more effective than NTP in learning non-local interactions; the use of transformer architectures with self-attention mechanisms; and the composition of the training dataset, which should include multispecies sequences to capture conserved regulatory signals across species and more accurately identify functional genomic elements.

Currently, NT models can process sequences up to only 6 kb. This poses a critical limitation for genomic applications, where the average gene length (exons + introns) spans *∼*27 kb, and key genes (e.g., CNTNAP2, ROBO2, NRXN3) exceed 1 Mb. To functionally assess genetic variants within whole-gene contexts, models must capture significantly longer-range dependencies, a capability hindered by the fundamental constraints of Transformer architectures.

The bottleneck stems from the self-attention mechanism, whose quadratic complexity (*O*(*n*^2^ *d*)) makes scaling to long sequences computationally intractable. For example, doubling the input length quadruples memory and compute demands, as attention requires pairwise token interactions. In practice, even sequences of modest length (e.g., 10-50 kb) become prohibitive due to the explosive growth of the *nxn* attention matrix.

Recent advances in transformer architectures have introduced innovative approaches to handle long sequences through modified attention mechanisms. Models employing sparse attention patterns, such as Longformer [16], or hashing-based approximations like Reformer [17], demonstrate how strategically constrained token interactions can maintain model performance while significantly reducing computational demands. Similarly, architectures incorporating memory mechanisms, including Transformer-XL [18], illustrate how intelligent sequence segmentation can extend context windows without incurring quadratic costs.

For genomic applications requiring analysis of megabase-scale sequences, there exists a compelling opportunity to explore these reduced-attention paradigms. While their computational advantages are well-established, their effectiveness in capturing complex biological relationships across extended genomic regions, such as long-range regulatory interactions or splicing patterns spanning large intronic regions, necessitates thorough investigation. The exploration should focus particularly on how different attention-sparsity patterns influence the model’s ability to maintain biological fidelity across various functional genomics tasks.

## Methods

### DNA foundation models

DNABERT was the first Transformer-based model specifically designed for DNA sequences analysis. DNABERT is a BERT-like model with 12 layers, 768 embedding dimensions, and 12 attention heads in each layer. It was trained with data generated from the human genome and takes as input sequences with a maximum length of 512 bp. DNABERT uses a tokenization strategy based on k-mers of 6 bp with overlap, allowing the model to capture local nucleotide dependencies effectively.

The NT was developed as a family of Transformer-based models with varying parameter sizes and training datasets. The NT models include versions trained on the human reference genome (GRCh38/hg38), incorporating genetic variants from the 1000 Genomes Project, as well as a model trained on all autosomal and sex chromosome sequences from GRCh38/hg38. Additionally, several NT models were trained on a large-scale multispecies dataset covering 850 species, including Bacteria, Fungi, Invertebrates, Protozoa, and Mammals. The NT models range in size from 2.5 billion to 50 million parameters, offering flexibility for different genomic analysis tasks. Unlike DNABERT, the NT models use a tokenization approach based on non-overlapping k-mers of 6 bases, allowing them to process longer sequences. NT models can analyze sequences up to 1000 tokens in length, corresponding to a total of 6 kilobase (kb).

HyenaDNA was introduced, comprising two models, HYENA 1 and HYENA_32. These models have 2 and 8 layers, respectively, both with 128 embedding dimensions and no attention heads. They were trained on the human reference genome and employ a distinct tokenization strategy based on single nucleotides, resulting in a vocabulary of only four genomic tokens (A, T, C, and G). Unlike DNABERT and NT, HyenaDNA models can process much longer sequences, with HYENA 1 handling up to 1000 bp and HYENA_32 extending to 32.000 bp, enabling efficient long-range genomic analysis. The Nucleotide Transformer models were obtained from https://github.com/instadeepai/nucleotide-transformer, the DNABERT from https://github.com/jerryji1993/DNABERT and the HyenaDNA models from https://github.com/HazyResearch/hyena-dna.

### CLINVAR

The ClinVar database in VCF format was downloaded from *ftp. ncbi. nlm. nih. gov/ pub/ clinvar/ vcf_ GRCh38/* . The VCF file was parsed to extract all benign SNVs (156.060) and pathogenic SNVs (65.865). Furthermore, we subsampled a set of 100.000 SNVs classified as uncertain significance, resulting in a total dataset of approximately 300.000 SNVs. For each variant, we extracted a segment of genomic sequence from the GRCh38/hg38 reference genome, centered on the ClinVar SNV. We then generated paired sequences for inference, with one sequence containing the reference base and the other incorporating the ClinVar variant. To fully leverage the capabilities of the DNA foundation models, we adjusted the sequence lengths based on their tokenization constraints. For the NT models, we used 999 tokens, with an additional Classification Token (CLS), resulting in a total of 1.000 tokens. DNABERT sequences were tokenized into 512 tokens, including the CLS token. HyenaDNA 1kb sequences consisted of 1.025 tokens, also incorporating the CLS token. For HyenaDNA 1kb and DNABERT, sequences were symmetrically trimmed to ensure that the Single nucleotide polymorphisms (SNP) was centrally positioned within the input window. In contrast, HyenaDNA 32kb, despite supporting a maximum input length of 32.770 tokens, processed the full 5.994-base sequence without requiring any modifications, as it fit within the model’s length constraints.

### Inference e Zero-shot scores

Each pair of sequences (the one containing the ClinVar variant and the other with the reference base) was processed by inference using each of the DNA foundation models. All inferences were conducted on an Nvidia A100 GPU Tensor Core with 40 GB of RAM, using custom scripts based on the PyTorch, Transformers, and Hugging Face libraries to efficiently handle model execution and data processing. For each sequence passed through the models, we extracted both the embeddings from the last layer and the logits. The logits were then transformed into probabilities using the softmax function.

To quantify the impact of each variant, we computed several distance metrics between the paired sequences. Specifically, for the embeddings, we calculated the Cosine, Manhattan, and Euclidean distances between corresponding token representations. For the probability distributions, we computed the

Hellinger, Jensen-Shannon, and Cross-Entropy distances between the output probabilities of each token pair. The Cumulative Context Score (CCS-N) is computed by aggregating Zero-shot scores at the variant site and its surrounding context in a distance-aware manner.

More precisely, let *s*_0_ denote the score at the central token (e.g., the token overlapping the variant), and let 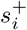 and 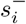 represent the scores at the *i*^*th*^ downstream and upstream tokens, respectively. The CCS-N is then defined as:

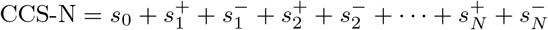

Scores are added symmetrically around the central token, expanding outward one pair at a time. This formulation enables the score to reflect increasingly broader contextual information while preserving sensitivity to positional proximity.

### U Statistic calculation

The Wilcoxon U-statistic measures the relative ranking of data points between two independent groups. Specifically, it quantifies how often a data point from one group ranks higher than a data point from the other, providing a non-parametric measure of distributional differences. In all our analyses, the Wilcoxon U-statistic was computed using the R function *wilcox*.*test*.

Since the number of SNVs differs across genetic consequence categories, we normalized the U-statistic by dividing it by its maximum possible value (*n*_1_ *× n*_2_, where *n*_1_ is the number of data points in the first group and *n*_2_ in the second) to allow meaningful comparisons.

To assess whether the models have learned the functional genomic structure surrounding each SNV, we annotated each token within the input sequences (6 kb for NT models, 512 bp for DNABERT, and bp for HYENA 1) into two distinct categories: coding regions (including first exons, internal exons, and last exons) and non-coding regions (including introns and intergenic regions). Annotations were based on GENCODE project annotation data (Release 47 for GRCh38, available at https://ftp.ebi.ac.uk/pub/databases/gencode/).

To evaluate whether different models captured the functional organization of the genome, we computed the Mann-Whitney U-statistic for each variant, comparing the Zero-shot scores of coding and non-coding tokens.

Since NT, DNABERT, and HyenaDNA models process different numbers of input tokens, we normalized the U-statistic by dividing it by its maximum possible value (*n*_1_ × *n*_2_, where *n*_1_ and *n*_2_ represent the number of coding and non-coding tokens, respectively), ensuring comparability across models.

