## Supplemental Figures for "Benchmarking DNA Foundation Models for zero-shot variant effect prediction: the role of context, training, and architecture"

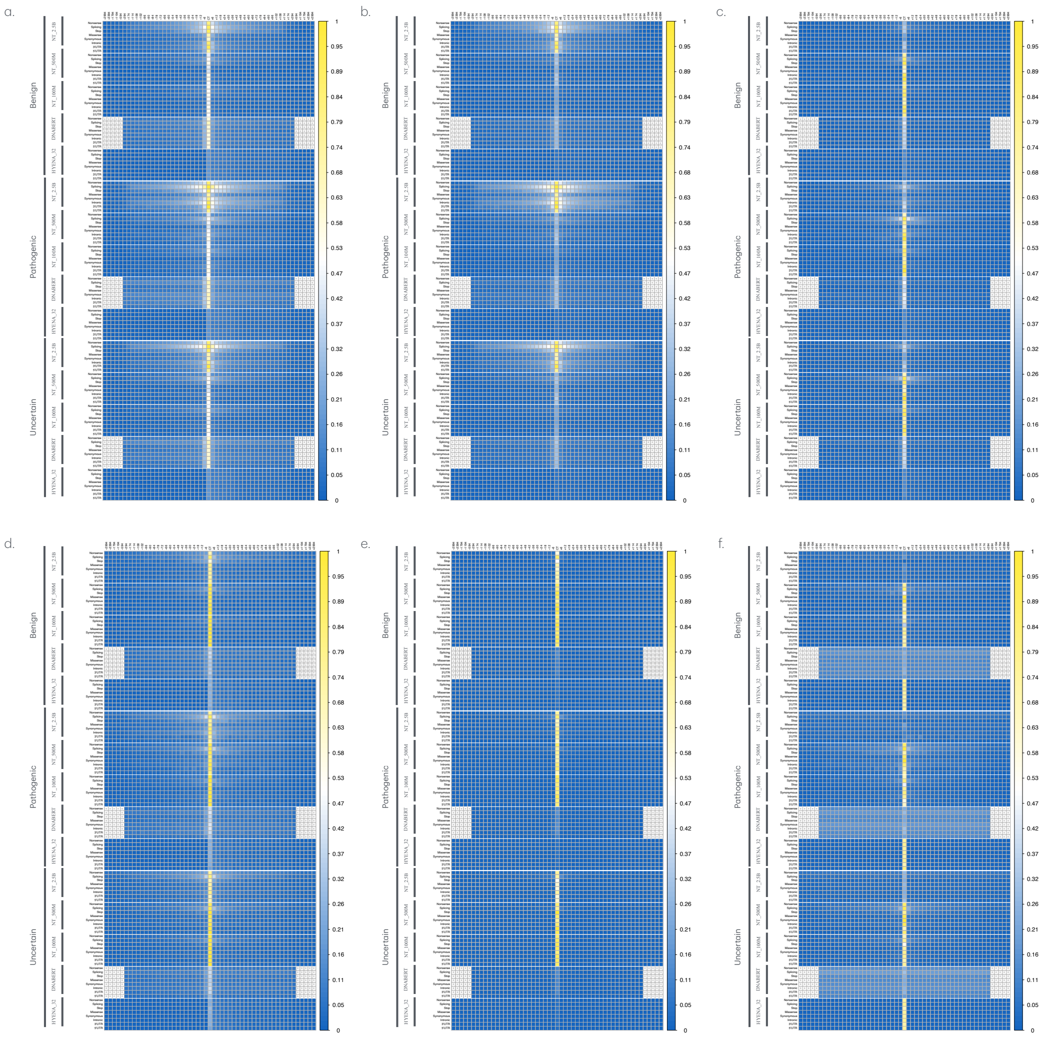


Figure S1: Figure displays the zero-shot score profiles across all tokens of the sequences analyzed using the DNA foundation models. Each row in the heatmap shows the trend of zero-shot score profiles calculated for sequence pairs (with ClinVar and reference bases) by averaging, for each token in the sequence, the zero-shot scores obtained by different models for each genetic consequence category (nonsense, stop, splicing, missense, synonymous, intronic, 5'UTR, and 3'UTR). The results are presented by distinguishing the three clinical impact categories (Benign, Pathogenic, and Uncertain) for the five main foundation models used (NT_2.5, NT_500, NT_100, DNABERT and HYENA_32). The intensity of each cell reflects the value of the zero-shot score according to the color legend on the right. Values are normalized by dividing each score by its maximum value. Panels (a-c) display zero-shot score profiles based on Euclidean (a), Manhattan (b), and Cosine (c) distances in the embedding space, while panels (d-f) show profiles based on Jensen-Shannon divergence (d), Cross-Entropy (e), and Hellinger distance (f) in the probability space.

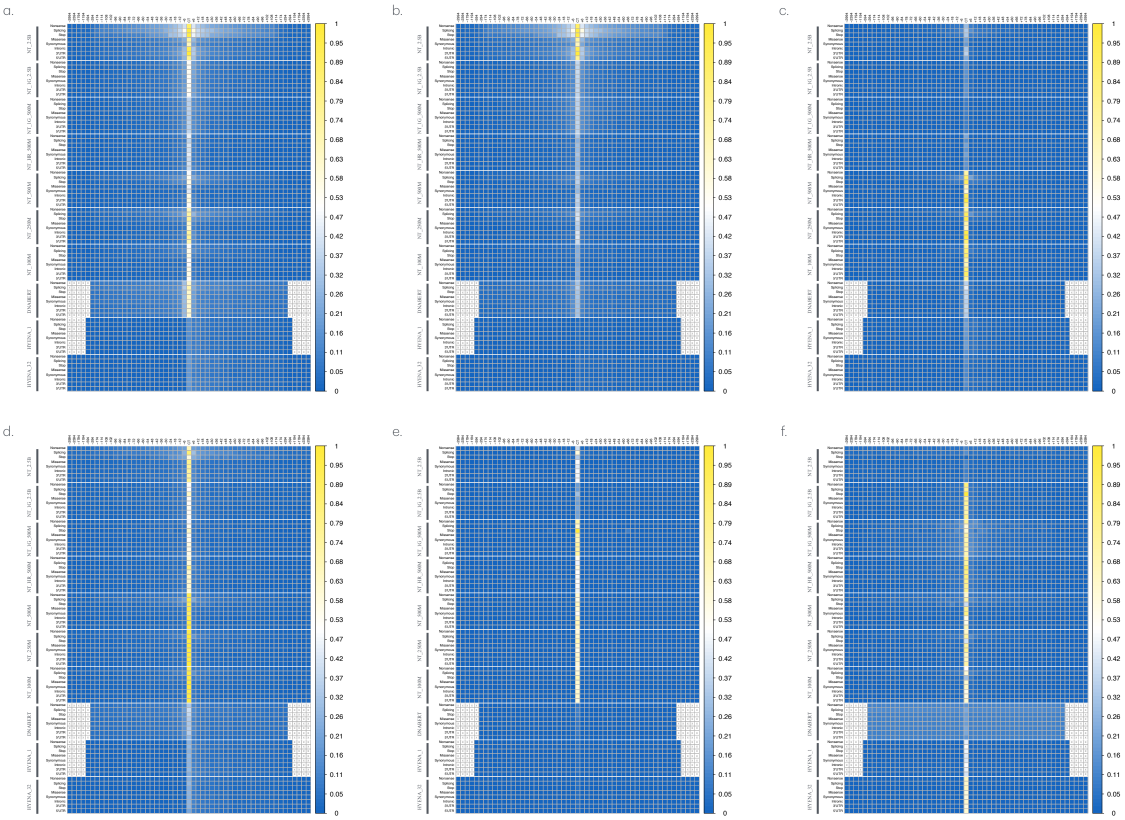

Figure S2: Figure displays the zero-shot score profiles of Clinvar Benign variants across all tokens of the sequences analyzed using the DNA foundation models. Each row in the heatmap shows the trend of zero-shot score profiles calculated for sequence pairs (with ClinVar and reference bases) by averaging, for each token in the sequence, the zero-shot scores obtained by different models for each genetic consequence category (nonsense, stop, splicing, missense, synonymous, intronic, 5'UTR, and 3'UTR). The results are presented for Clinvar Benign variants for all the ten foundation models used (NT_2.5, NT_1G_2.5B, NT_1G_500M, NT_HR_500M, NT_500, NT_250, NT_100, DNABERT, HYENA_1 and HYENA_32). The intensity of each cell reflects the value of the zero-shot score according to the color legend on the right. Values are normalized by dividing each score by its maximum value. Panels (a–c) display zero-shot score profiles based on Euclidean (a), Manhattan (b), and Cosine (c) distances in the embedding space, while panels (d–f) show profiles based on Jensen-Shannon divergence (d), Cross-Entropy (e), and Hellinger distance (f) in the probability space.

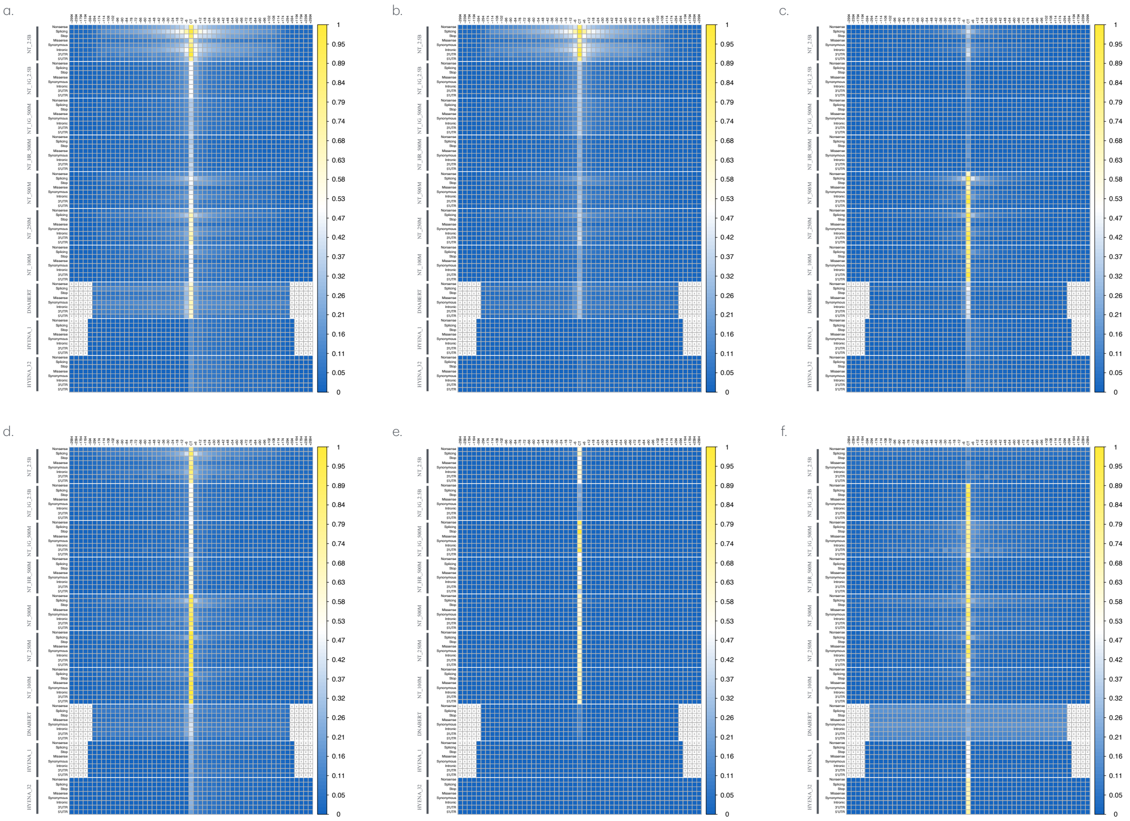

Figure S3: Figure displays the zero-shot score profiles of Clinvar Pathogenic variants across all tokens of the sequences analyzed using the DNA foundation models. Each row in the heatmap shows the trend of zero-shot score profiles calculated for sequence pairs (with ClinVar and reference bases) by averaging, for each token in the sequence, the zero-shot scores obtained by different models for each genetic consequence category (nonsense, stop, splicing, missense, synonymous, intronic, 5'UTR, and 3'UTR). The results are presented for Clinvar Pathogenic variants for all the ten foundation models used (NT_2.5, NT_1G_2.5B, NT_1G_500M, NT_HR_500M, NT_500, NT_250, NT_100, DNABERT, HYENA_1 and HYENA_32). The intensity of each cell reflects the value of the zero-shot score according to the color legend on the right. Values are normalized by dividing each score by its maximum value. Panels (a–c) display zero-shot score profiles based on Euclidean (a), Manhattan (b), and Cosine (c) distances in the embedding space, while panels (d–f) show profiles based on Jensen-Shannon divergence (d), Cross-Entropy (e), and Hellinger distance (f) in the probability space.

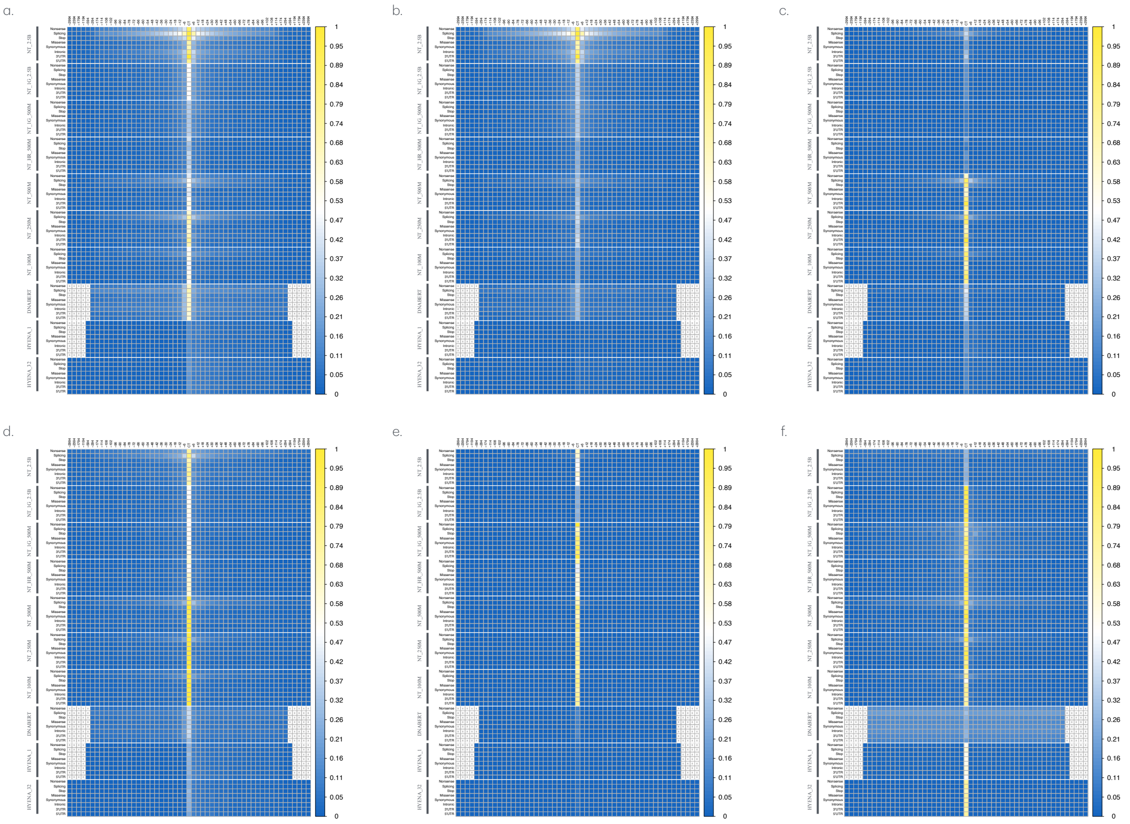


Figure S4: Figure displays the zero-shot score profiles of Clinvar Uncertain Significance variants across all tokens of the sequences analyzed using the DNA foundation models. Each row in the heatmap shows the trend of zero-shot score profiles calculated for sequence pairs (with ClinVar and reference bases) by averaging, for each token in the sequence, the zero-shot scores obtained by different models for each genetic consequence category (nonsense, stop, splicing, missense, synonymous, intronic, 5'UTR, and 3'UTR). The results are presented for Clinvar Pathogenic variants for all the ten foundation models used (NT_2.5, NT_1G_2.5B, NT_1G_500M, NT_HR_500M, NT_500, NT_250, NT_100, DNABERT, HYENA_1 and HYENA_32). The intensity of each cell reflects the value of the zero-shot score according to the color legend on the right. Values are normalized by dividing each score by its maximum value. Panels (a-c) display zero-shot score profiles based on Euclidean (a), Manhattan (b), and Cosine (c) distances in the embedding space, while panels (d-f) show profiles based on Jensen-Shannon divergence (d), Cross-Entropy (e), and Hellinger distance (f) in the probability space.

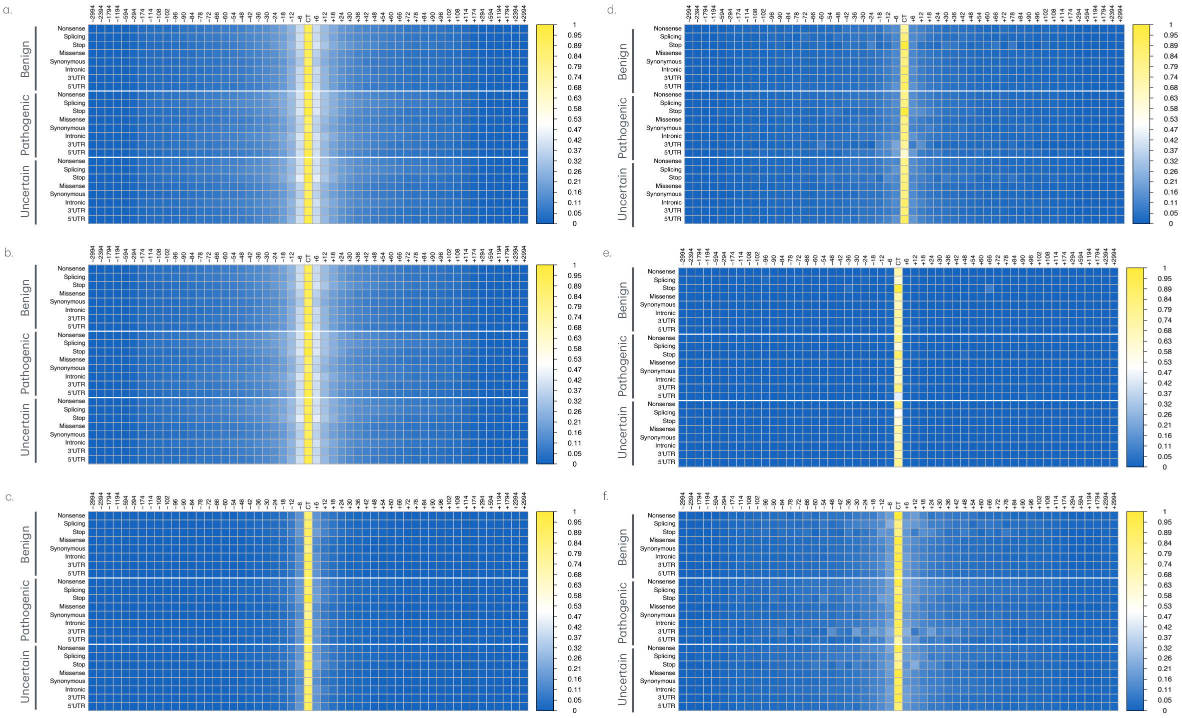


Figure S5: Figure displays the zero-shot score profiles across all tokens of the sequences analyzed using the NT_1G_2.5B foundation models. Each row in the heatmap shows the trend of zero-shot score profiles calculated for sequence pairs (with ClinVar and reference bases) by averaging, for each token in the sequence, the zero-shot scores obtained by different models for each genetic consequence category (nonsense, stop, splicing, missense, synonymous, intronic, 5'UTR, and 3'UTR). The results are presented by distinguishing the three clinical impact categories (Benign, Pathogenic, and Uncertain) for the NT_1G_2.5B model . The intensity of each cell reflects the value of the zero-shot score according to the color legend on the right. Values are normalized by dividing each score by its maximum value. Panels (a-c) display zero-shot score profiles based on Euclidean (a), Manhattan (b), and Cosine (c) distances in the embedding space, while panels (d-f) show profiles based on Jensen-Shannon divergence (d), Cross-Entropy (e), and Hellinger distance (f) in the probability space.

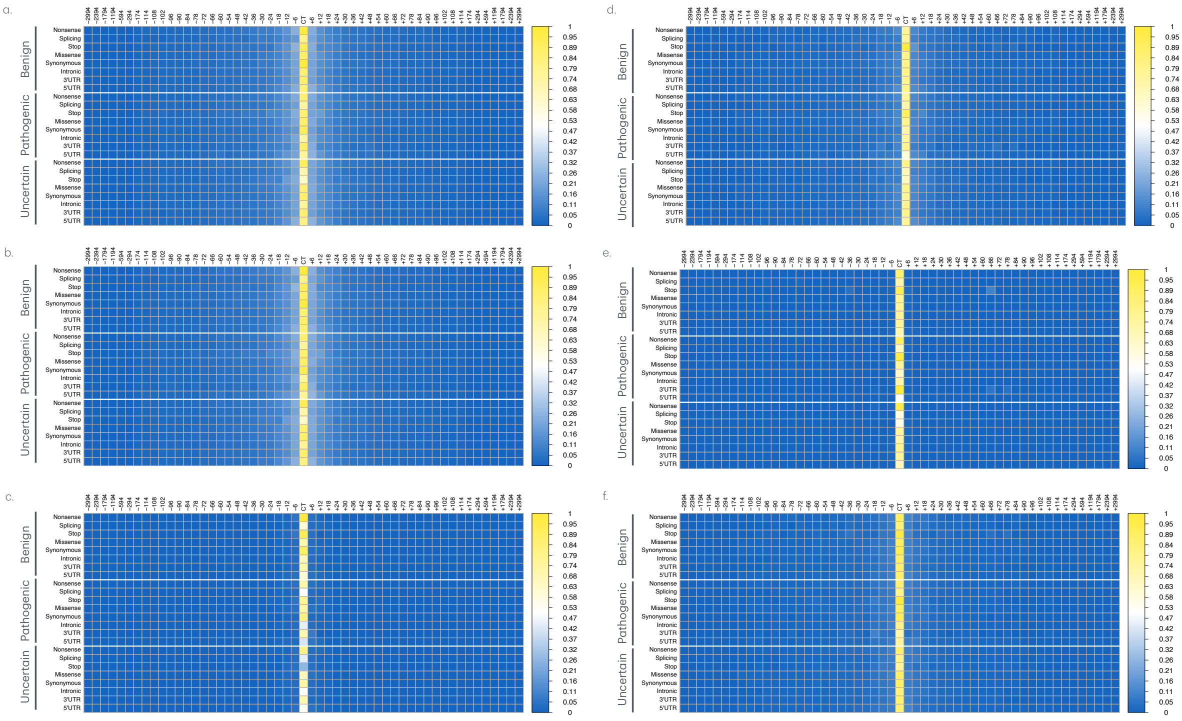


Figure S6: Figure displays the zero-shot score profiles across all tokens of the sequences analyzed using the NT_1G_500M foundation models. Each row in the heatmap shows the trend of zero-shot score profiles calculated for sequence pairs (with ClinVar and reference bases) by averaging, for each token in the sequence, the zero-shot scores obtained by different models for each genetic consequence category (nonsense, stop, splicing, missense, synonymous, intronic, 5'UTR, and 3'UTR). The results are presented by distinguishing the three clinical impact categories (Benign, Pathogenic, and Uncertain) for the NT_1G_500M model . The intensity of each cell reflects the value of the zero-shot score according to the color legend on the right. Values are normalized by dividing each score by its maximum value. Panels (a-c) display zero-shot score profiles based on Euclidean (a), Manhattan (b), and Cosine (c) distances in the embedding space, while panels (d-f) show profiles based on Jensen-Shannon divergence (d), Cross-Entropy (e), and Hellinger distance (f) in the probability space.

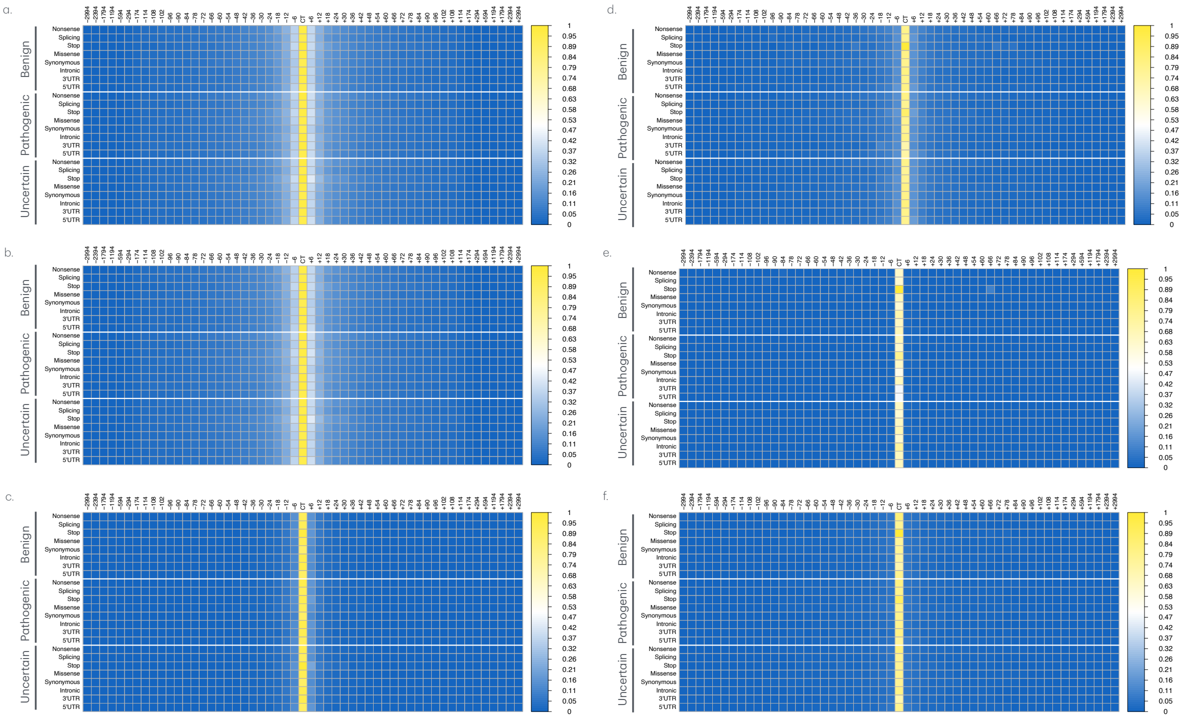


Figure S7: Figure displays the zero-shot score profiles across all tokens of the sequences analyzed using the NT_HR foundation models. Each row in the heatmap shows the trend of zero-shot score profiles calculated for sequence pairs (with ClinVar and reference bases) by averaging, for each token in the sequence, the zero-shot scores obtained by different models for each genetic consequence category (nonsense, stop, splicing, missense, synonymous, intronic, 5'UTR, and 3'UTR). The results are presented by distinguishing the three clinical impact categories (Benign, Pathogenic, and Uncertain) for the NT_HR model. The intensity of each cell reflects the value of the zero-shot score according to the color legend on the right. Values are normalized by dividing each score by its maximum value. Panels (a-c) display zero-shot score profiles based on Euclidean (a), Manhattan (b), and Cosine (c) distances in the embedding space, while panels (d-f) show profiles based on Jensen-Shannon divergence (d), Cross-Entropy (e), and Hellinger distance (f) in the probability space.

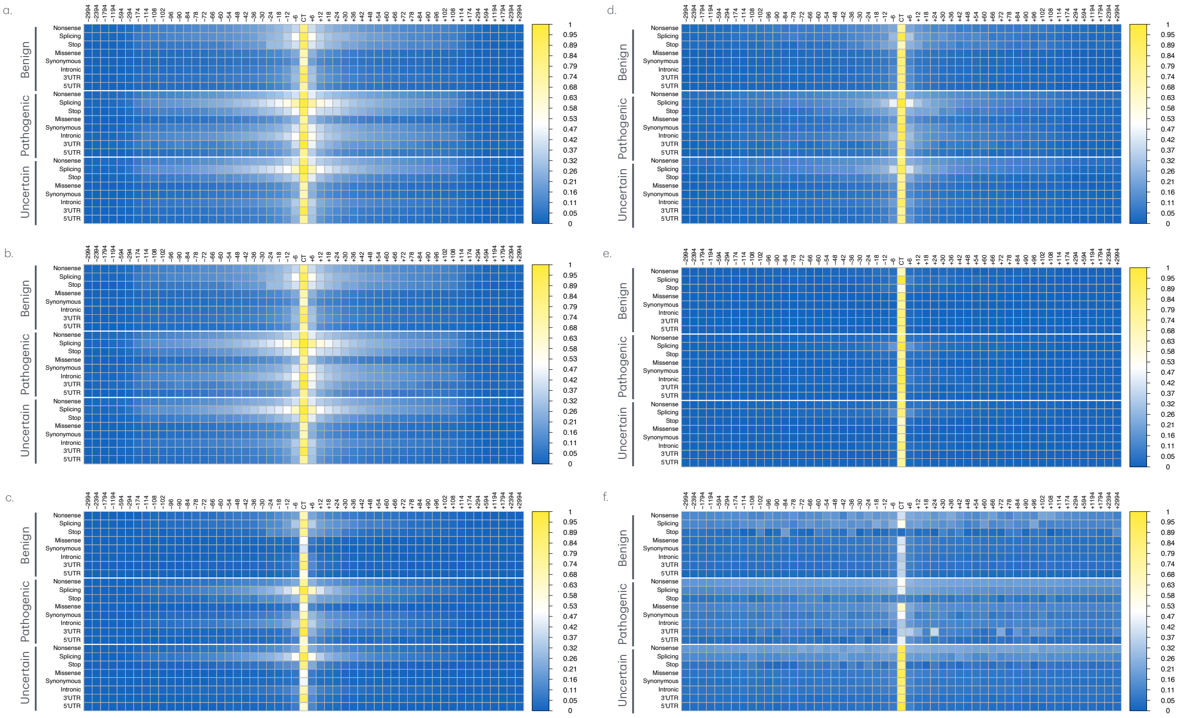

Figure S8: Figure displays the zero-shot score profiles across all tokens of the sequences analyzed using the NT_2.5B foundation models. Each row in the heatmap shows the trend of zero-shot score profiles calculated for sequence pairs (with ClinVar and reference bases) by averaging, for each token in the sequence, the zero-shot scores obtained by different models for each genetic consequence category (nonsense, stop, splicing, missense, synonymous, intronic, 5'UTR, and 3'UTR). The results are presented by distinguishing the three clinical impact categories (Benign, Pathogenic, and Uncertain) for the NT_2.5B model . The intensity of each cell reflects the value of the zero-shot score according to the color legend on the right. Values are normalized by dividing each score by its maximum value. Panels (a-c) display zero-shot score profiles based on Euclidean (a), Manhattan (b), and Cosine (c) distances in the embedding space, while panels (d-f) show profiles based on Jensen-Shannon divergence (d), Cross-Entropy (e), and Hellinger distance (f) in the probability space.

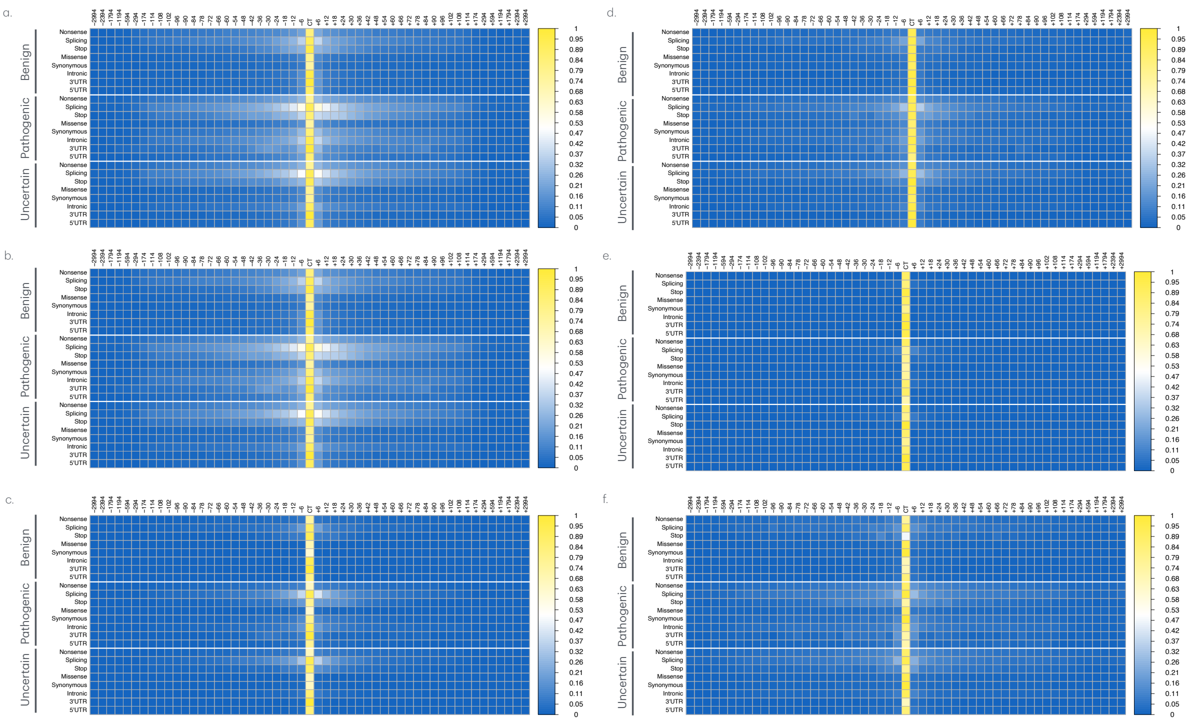


Figure S9: Figure displays the zero-shot score profiles across all tokens of the sequences analyzed using the NT_500M foundation models. Each row in the heatmap shows the trend of zero-shot score profiles calculated for sequence pairs (with ClinVar and reference bases) by averaging, for each token in the sequence, the zero-shot scores obtained by different models for each genetic consequence category (nonsense, stop, splicing, missense, synonymous, intronic, 5'UTR, and 3'UTR). The results are presented by distinguishing the three clinical impact categories (Benign, Pathogenic, and Uncertain) for the NT_500M model . The intensity of each cell reflects the value of the zero-shot score according to the color legend on the right. Values are normalized by dividing each score by its maximum value. Panels (a-c) display zero-shot score profiles based on Euclidean (a), Manhattan (b), and Cosine (c) distances in the embedding space, while panels (d-f) show profiles based on Jensen-Shannon divergence (d), Cross-Entropy (e), and Hellinger distance (f) in the probability space.

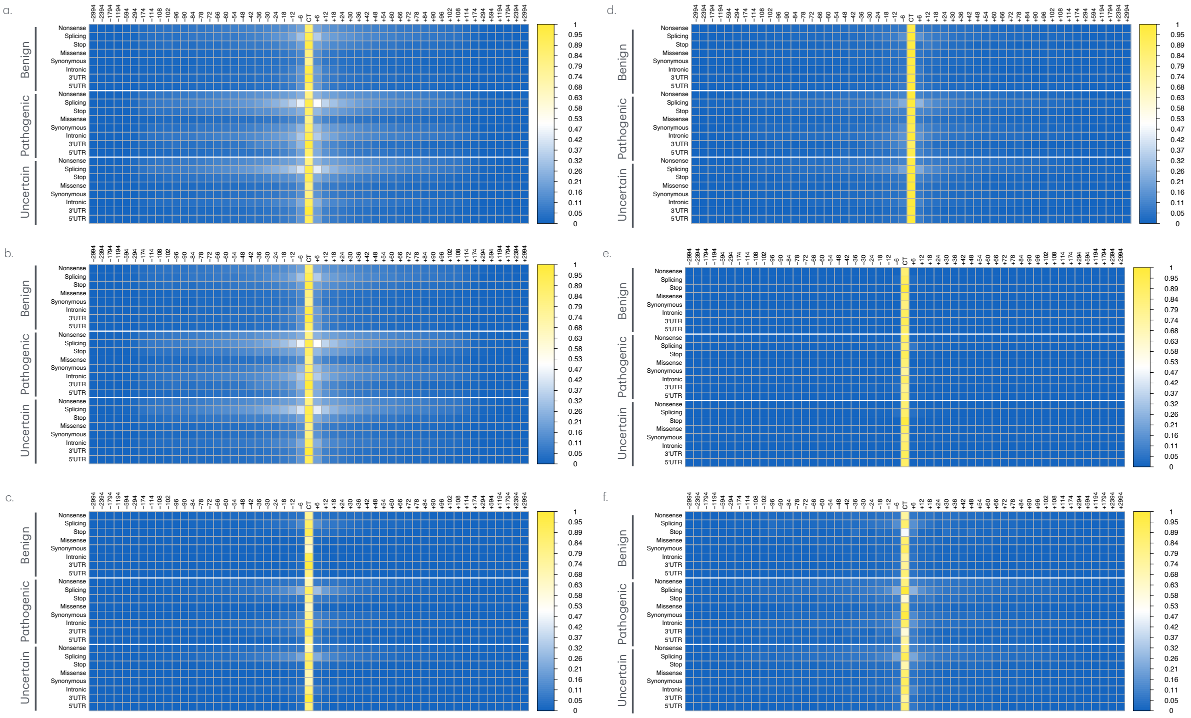


Figure S10: Figure displays the zero-shot score profiles across all tokens of the sequences analyzed using the NT_250M foundation models. Each row in the heatmap shows the trend of zero-shot score profiles calculated for sequence pairs (with ClinVar and reference bases) by averaging, for each token in the sequence, the zero-shot scores obtained by different models for each genetic consequence category (nonsense, stop, splicing, missense, synonymous, intronic, 5'UTR, and 3'UTR). The results are presented by distinguishing the three clinical impact categories (Benign, Pathogenic, and Uncertain) for the NT_250M model . The intensity of each cell reflects the value of the zero-shot score according to the color legend on the right. Values are normalized by dividing each score by its maximum value. Panels (a-c) display zero-shot score profiles based on Euclidean (a), Manhattan (b), and Cosine (c) distances in the embedding space, while panels (d-f) show profiles based on Jensen-Shannon divergence (d), Cross-Entropy (e), and Hellinger distance (f) in the probability space.

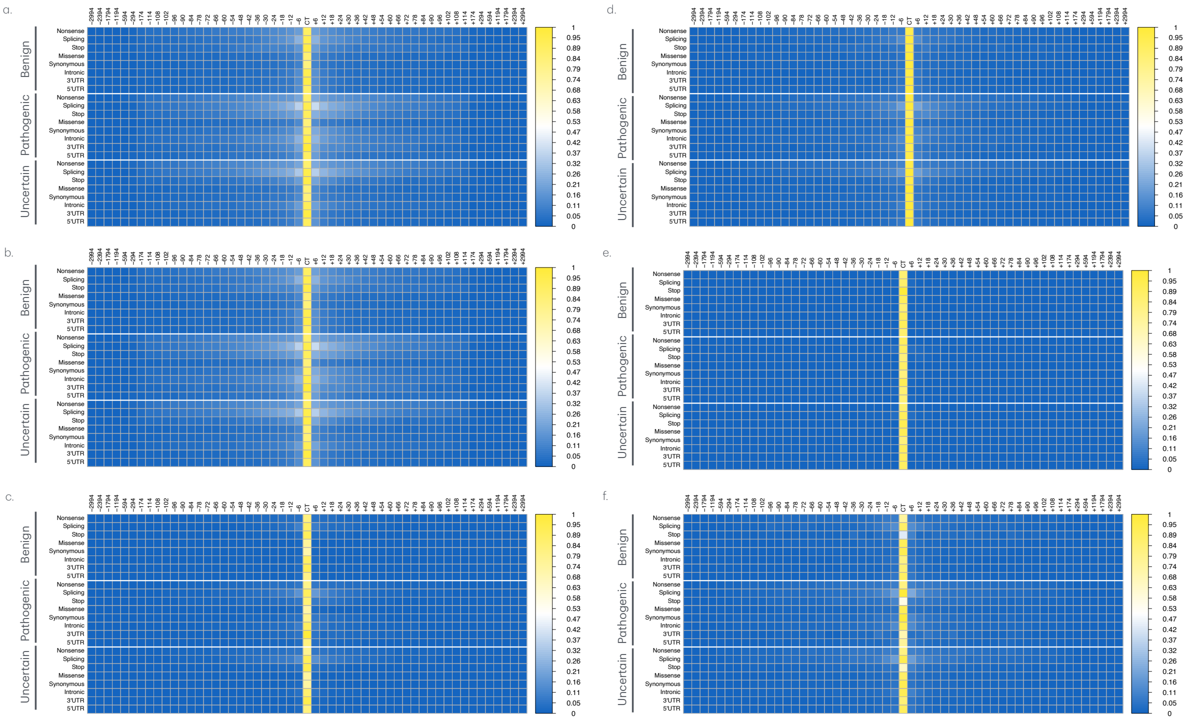


Figure S11: Figure displays the zero-shot score profiles across all tokens of the sequences analyzed using the NT_100M foundation models. Each row in the heatmap shows the trend of zero-shot score profiles calculated for sequence pairs (with ClinVar and reference bases) by averaging, for each token in the sequence, the zero-shot scores obtained by different models for each genetic consequence category (nonsense, stop, splicing, missense, synonymous, intronic, 5'UTR, and 3'UTR). The results are presented by distinguishing the three clinical impact categories (Benign, Pathogenic, and Uncertain) for the NT_100M model . The intensity of each cell reflects the value of the zero-shot score according to the color legend on the right. Values are normalized by dividing each score by its maximum value. Panels (a-c) display zero-shot score profiles based on Euclidean (a), Manhattan (b), and Cosine (c) distances in the embedding space, while panels (d-f) show profiles based on Jensen-Shannon divergence (d), Cross-Entropy (e), and Hellinger distance (f) in the probability space.

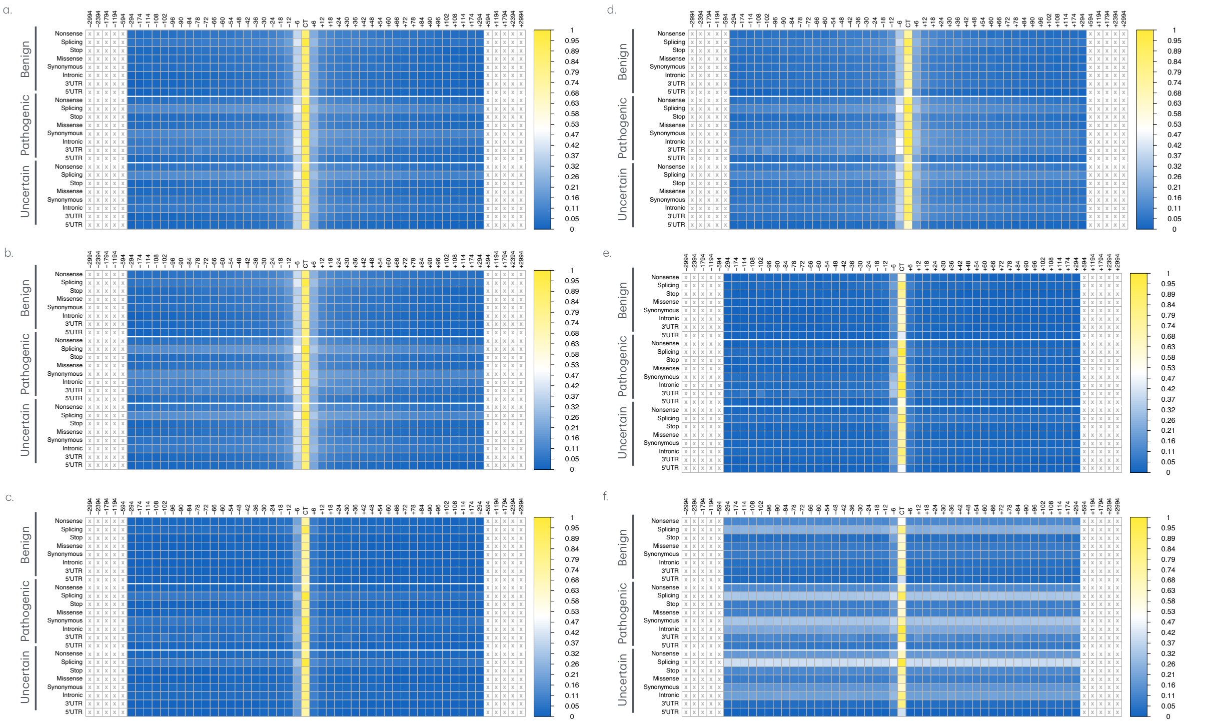


Figure S12: Figure displays the zero-shot score profiles across all tokens of the sequences analyzed using the DNABERT foundation models. Each row in the heatmap shows the trend of zero-shot score profiles calculated for sequence pairs (with ClinVar and reference bases) by averaging, for each token in the sequence, the zero-shot scores obtained by different models for each genetic consequence category (nonsense, stop, splicing, missense, synonymous, intronic, 5'UTR, and 3'UTR). The results are presented by distinguishing the three clinical impact categories (Benign, Pathogenic, and Uncertain) for the DNABERT model . The intensity of each cell reflects the value of the zero-shot score according to the color legend on the right. Values are normalized by dividing each score by its maximum value. Panels (a-c) display zero-shot score profiles based on Euclidean (a), Manhattan (b), and Cosine (c) distances in the embedding space, while panels (d-f) show profiles based on Jensen-Shannon divergence (d), Cross-Entropy (e), and Hellinger distance (f) in the probability space.

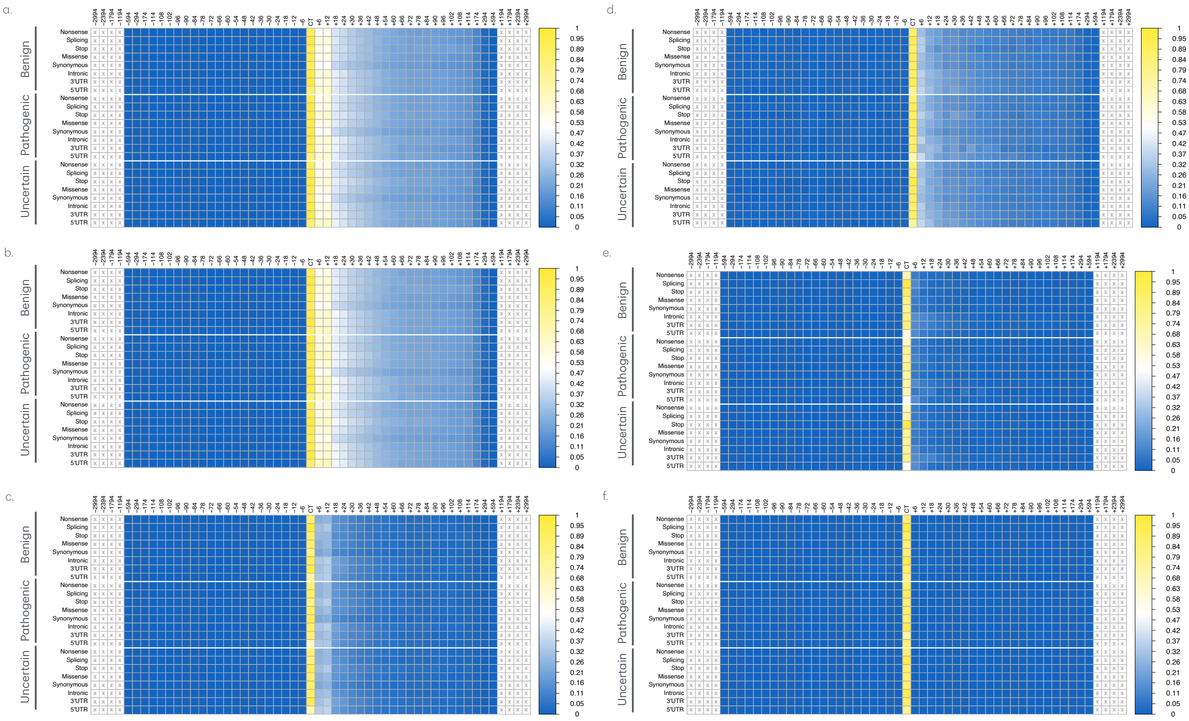


Figure S13: Figure displays the zero-shot score profiles across all tokens of the sequences analyzed using the Hyena_1 foundation models. Each row in the heatmap shows the trend of zero-shot score profiles calculated for sequence pairs (with ClinVar and reference bases) by averaging, for each token in the sequence, the zero-shot scores obtained by different models for each genetic consequence category (nonsense, stop, splicing, missense, synonymous, intronic, 5'UTR, and 3'UTR). The results are presented by distinguishing the three clinical impact categories (Benign, Pathogenic, and Uncertain) for theHyena_1 model . The intensity of each cell reflects the value of the zero-shot score according to the color legend on the right. Values are normalized by dividing each score by its maximum value. Panels (a-c) display zero-shot score profiles based on Euclidean (a), Manhattan (b), and Cosine (c) distances in the embedding space, while panels (d-f) show profiles based on Jensen-Shannon divergence (d), Cross-Entropy (e), and Hellinger distance (f) in the probability space.

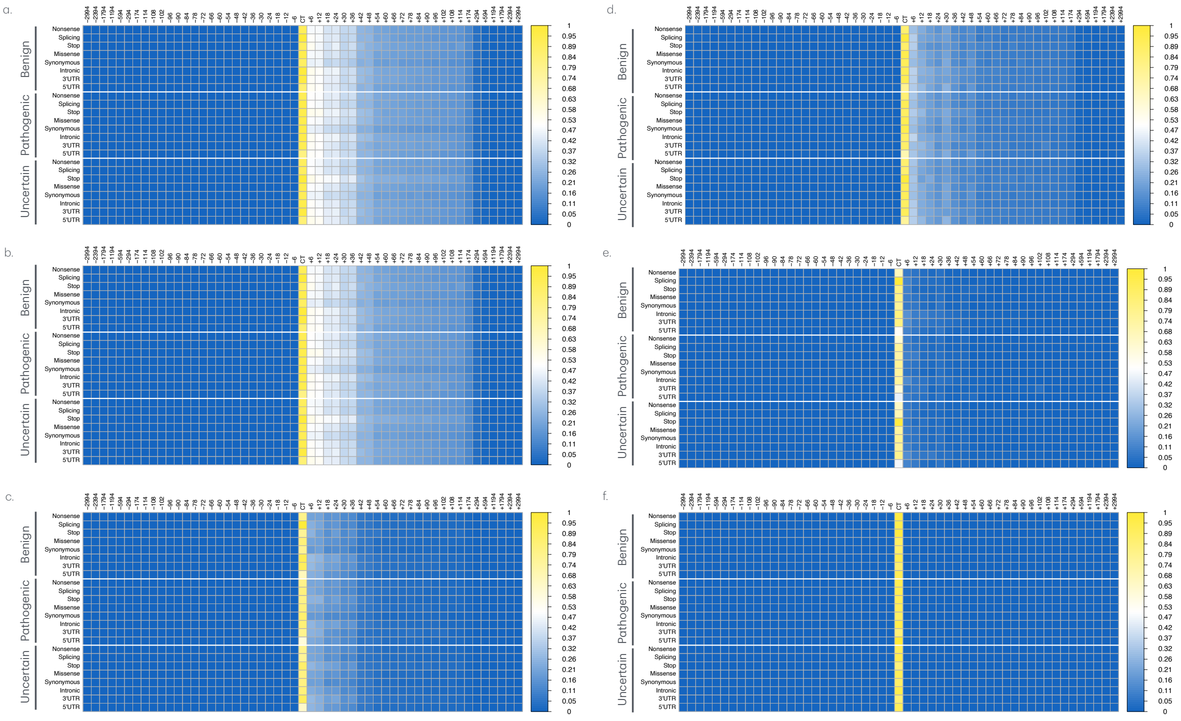


Figure S14: Figure displays the zero-shot score profiles across all tokens of the sequences analyzed using the Hyena_32 foundation models. Each row in the heatmap shows the trend of zero-shot score profiles calculated for sequence pairs (with ClinVar and reference bases) by averaging, for each token in the sequence, the zero-shot scores obtained by different models for each genetic consequence category (nonsense, stop, splicing, missense, synonymous, intronic, 5'UTR, and 3'UTR). The results are presented by distinguishing the three clinical impact categories (Benign, Pathogenic, and Uncertain) for the Hyena_32 model . The intensity of each cell reflects the value of the zero-shot score according to the color legend on the right. Values are normalized by dividing each score by its maximum value. Panels (a-c) display zero-shot score profiles based on Euclidean (a), Manhattan (b), and Cosine (c) distances in the embedding space, while panels (d-f) show profiles based on Jensen-Shannon divergence (d), Cross-Entropy (e), and Hellinger distance (f) in the probability space.

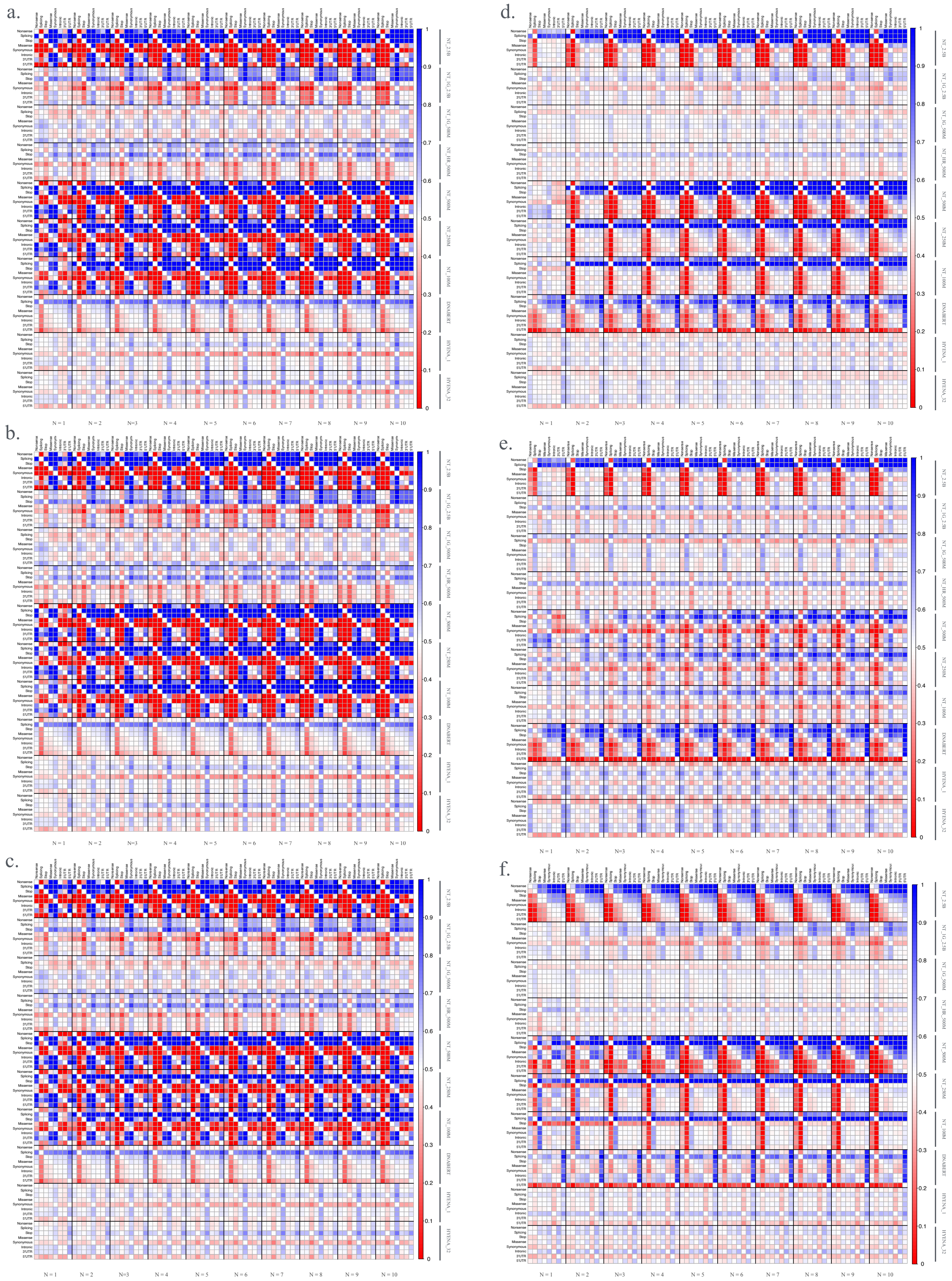


Figure S15: The figure illustrates the ability of the ten DNA foundation models to distinguish benign from pathogenic variants. Each point in the dot chart represents the normalized Wilcoxon U-statistic, comparing the distribution of CCS-N, obtained by summing the zero-shot scores of the N tokens closest to the central token carrying the ClinVar variants, between benign and pathogenic variants within the same genetic consequence class. Panels (a-c) shows the results for Euclidean (a), Manhattan (b), and Cosine (c) distances in the embedding space, while panels (d-f) show results based on Jensen-Shannon divergence (d), Cross-Entropy (e), and Hellinger distance (f) in the probability space.

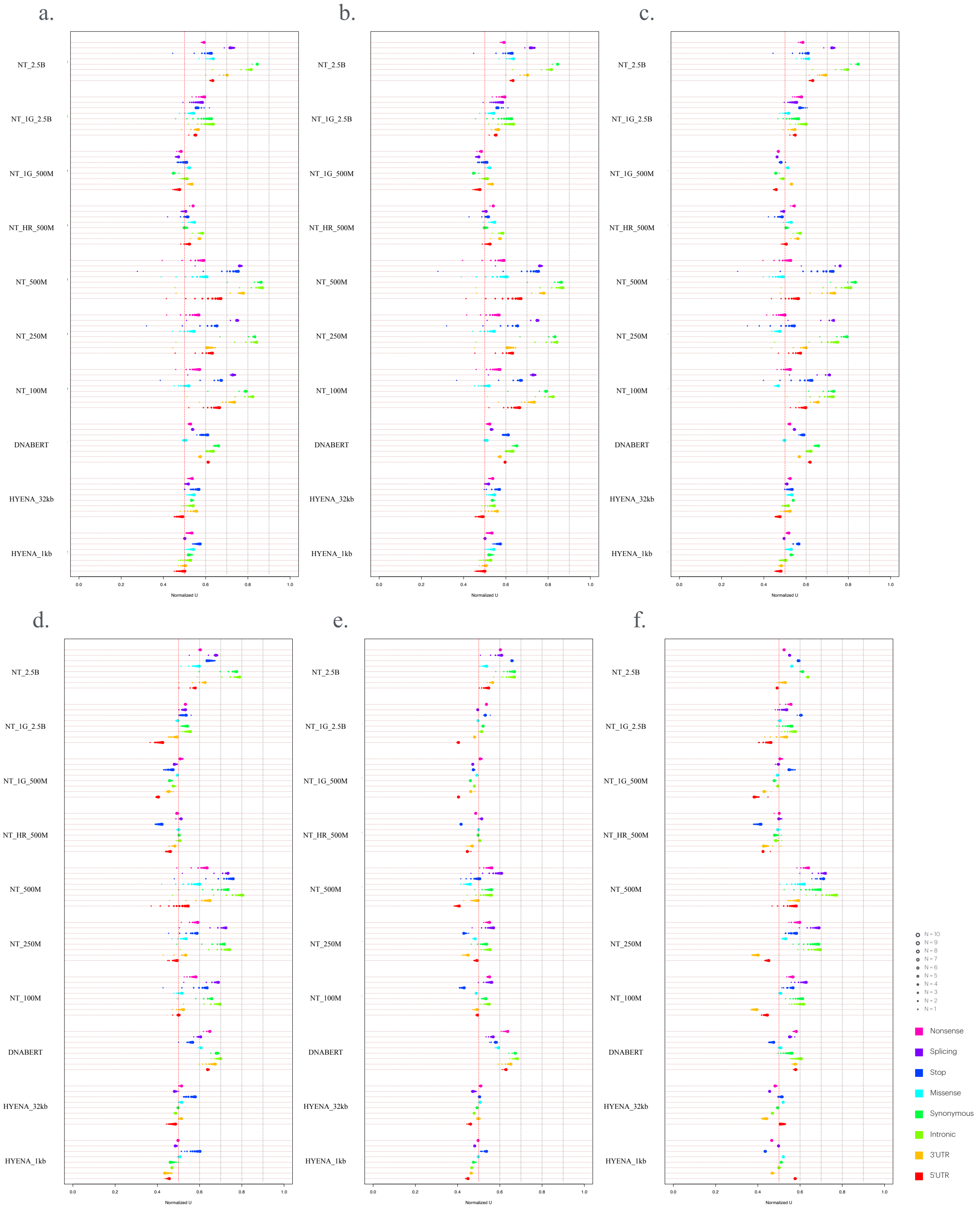

Figure S16: The figure illustrates the ability of the ten DNA foundation models to distinguish benign from pathogenic variants. Each point in the dot chart represents the normalized Wilcoxon U-statistic, comparing the distribution of CCS-N, obtained by summing the zero-shot scores of the N tokens closest to the central token carrying the ClinVar variants, between benign and pathogenic variants within the same genetic consequence class. Panels (a-c) shows the results for Euclidean (a), Manhattan (b), and Cosine (c) distances in the embedding space, while panels (d-f) show results based on Jensen-Shannon divergence (d), Cross-Entropy (e), and Hellinger distance (f) in the probability space.

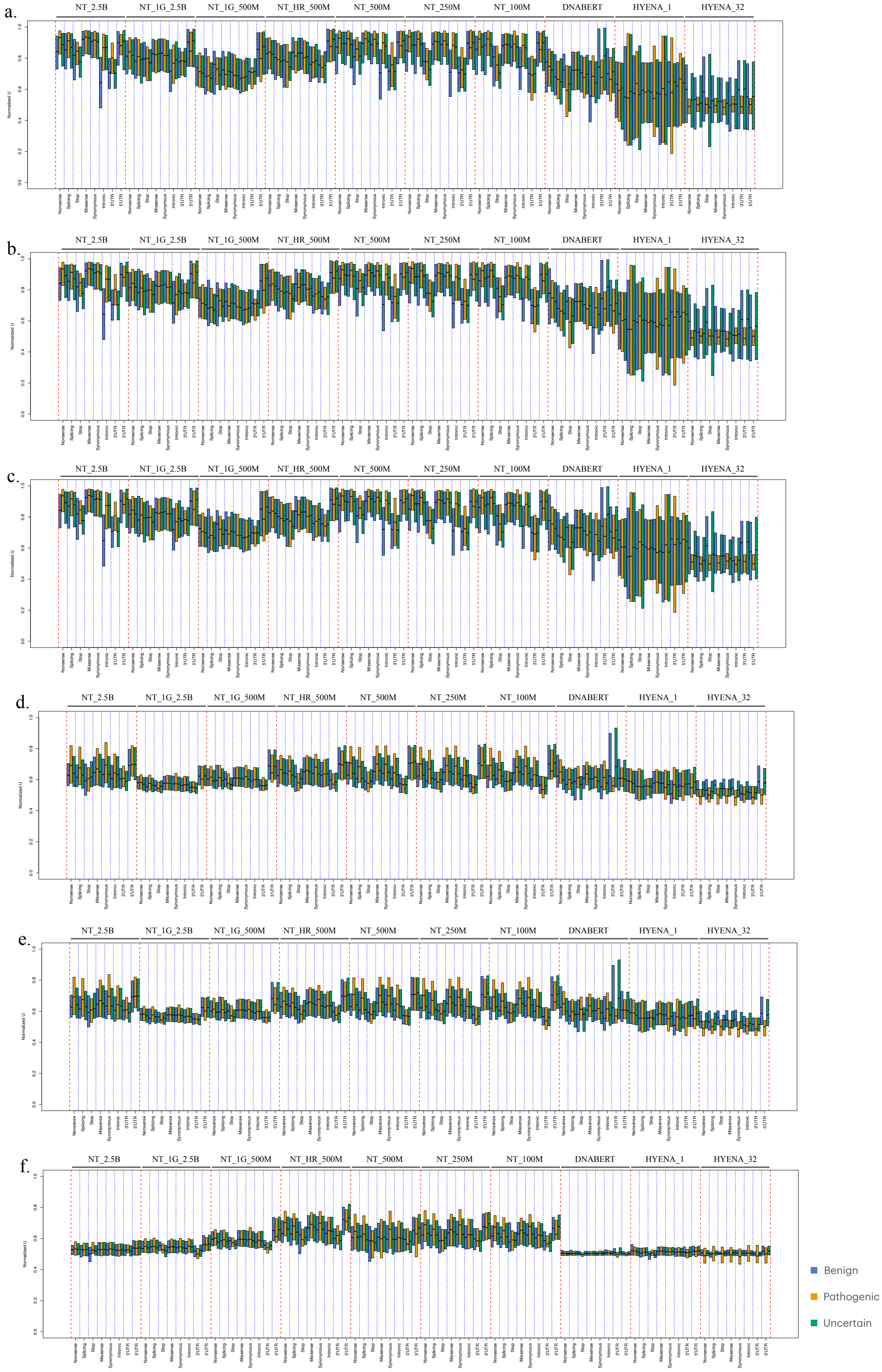


Figure S17: The figure illustrates the ability of the ten foundation models to assess the impact of ClinVar variants on the functional structure of the genome. The box plots show, for each genetic consequence class, the distribution of normalized U-statistics comparing zero-shot scores between coding elements (exons) and non-coding elements (introns). Panels (a-c) shows the results for Euclidean (a), Manhattan (b), and Cosine (c) distances in the embedding space, while panels (d-f) show results based on Jensen-Shannon divergence (d), Cross-Entropy (e), and Hellinger distance (f) in the probability space.
